# A moving front of osteoblast maturation scales regenerating zebrafish bone

**DOI:** 10.64898/2026.09.14.745769

**Authors:** Konrad Marx, Simone Cicolini, Delphine Colevret, Coline Coudeville, Guillaume Salbreux, Alessandro De Simone

## Abstract

Successful regeneration recovers lost tissues to their original form. Here, we investigate how proliferation of bone-forming osteoblasts is timely regulated to control tissue size during zebrafish scale regeneration. Regeneration of the scale osteoblast tissue proceeds in two phases: an initial osteoblast proliferative expansion followed by hypertrophic growth without cell division. First, we show that regeneration recovers scale size by amplifying the number of osteoblasts by a fixed factor, starting from a properly scaled tissue primordium. This fixed expansion is achieved by regulating proliferation such that osteoblast number increases roughly linearly, with the rate of increase scaling with initial osteoblast number, and halting cell proliferation at a given time. To understand the mechanisms underlying this proliferation control, we used live imaging of transgenic reporters and biosensors together with transcriptomics. We uncover that proliferation arrest is imparted by a moving front of osteoblast maturation travelling outward across the scale, with dynamics that scales with tissue size. The osteoblast maturation front coincides with onset of hypertrophy and bone deposition, and it is accompanied by reduction in expression of the *fibroblast growth factor receptor 3* (*fgfr3*) and activity of the kinases Erk (Extracellular signal-Regulated Kinase) and p38. Inhibition of Fgf/Erk signaling and osteoblast proliferation does not alter the progression of the maturation front. We show that the scaling maturation front can quantitatively explain the osteoblast proliferation dynamics and its scaling across different tissue sizes. Overall, we identify a moving front of cell maturation as a mechanism for controlling cell proliferation across a regenerating tissue.

## Introduction

In regeneration, a lost or damaged body part is restored to its original form. Regeneration requires the tight coordination of cellular events, such as differentiation, proliferation and growth (*1–4*). These processes must be coordinated across large distances and arrested in a timely manner for regenerating tissues to achieve their appropriate size and shape. Several models have been proposed to explain proliferation control in developmental and regenerative contexts (*5, 6*). In the “division counter” model, cells proliferate for a certain number of cell cycles, then stop. Alternatively, in the “timer” model, cells proliferate for a given time. Counters and timers could in principle be cell-autonomous or emergent properties of the cell population. Among the models acting at the population level, in the “sizer” model, a mechanism provides cells with information on tissue growth or current size, such that proliferation is modulated accordingly and arrested once a target tissue size is reached. In many regenerating systems, such target size is the size of the tissue before injury, and it can vary between body locations and animals. Therefore, counters, timers and sizers need to be tuned to recover a variable original size.

In most regenerating systems, it remains unclear which, if any, of these mechanisms would act and what molecular and/or physical cues would measure the total number of cell divisions, time or tissue size. In addition, it is unclear how this information would modulate cell proliferation across a tissue to achieve the correct final number of cells. Several key cellular and molecular events that are important for controlling proliferation in regenerating tissues have been described (*1–4*), but the study of their coordination has been limited by the inaccessibility of these tissues to real-time imaging (*7*). Here, we investigate the control of osteoblast proliferation in regenerating zebrafish scales, external dermal appendages accessible to live microscopy (*8–13*).

Scales are dermal bone disks that each consist of a central layer of bone matrix covered by a monolayer of bone-forming cells (Figure 1A) (*14, 15*), referred to as “osteoblasts” (*8–10, 16–18*). After a scale is lost, osteoblast regeneration proceeds through a stereotypical sequence of events (*9*). First, a new osteoblast tissue primordium forms from the surrounding tissue. This newly formed osteoblast pool proliferates for a few days (Figure 1B, proliferation). Osteoblast number increases until it reaches a plateau about 5 days after the onset of regeneration (days post-plucking; dpp) (*9*)). Then, proliferation stops in the bulk of the scale and cells continue growing by hypertrophy, i.e. cell growth without division (Figure 1B, hypertrophy). The progression from differentiation from a mesenchymal progenitor to an early proliferative stage and then to mature bone-forming and mineralizing stages is a shared feature of osteoblasts in many systems, including in zebrafish fins (*19–24*). This progression is accompanied by the expression of characteristic genes, such as the transcription factors *runx2* and *osterix (osx* or *sp7),* and bone formation genes, such as *osteonectin* (*17–21*). At the same time, Mitogen-Activated Pathway Kinases (MAPKs) such as the Extracellular signal-Regulated Kinase (Erk) and p38 are essential for osteoblast proliferation and differentiation in development and regeneration (*25–28*). However, it is not clear how these processes are spatially organized and timed to control tissue size and architecture.

**Figure 1.**
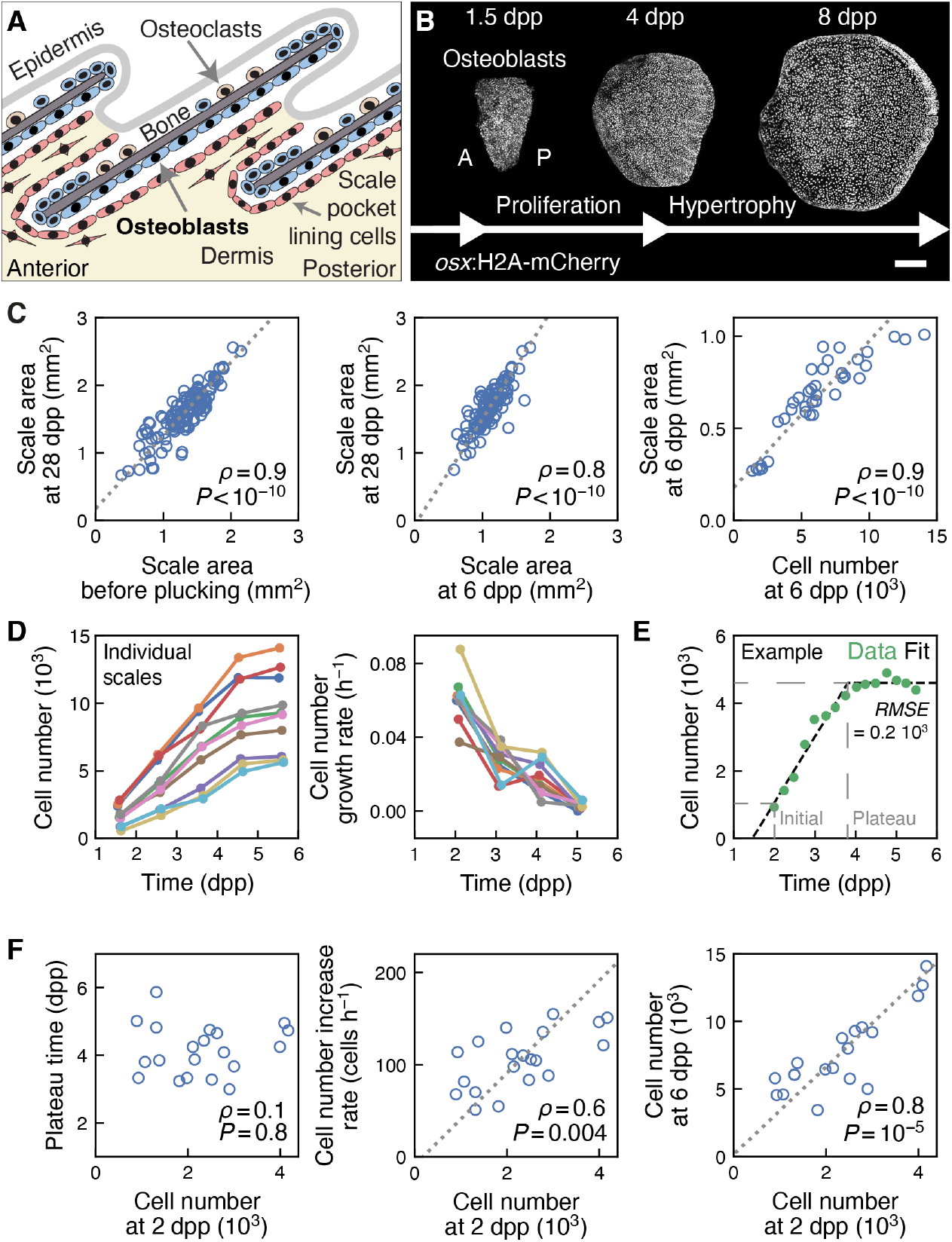
Dynamics and scaling of scale osteoblast proliferation. **A**. Schematics of zebrafish scale morphology, depicted in cross-section. **B**. Phases of scale regeneration with examples of scale osteoblasts. **C**. Left: area of each regenerated scale compared to the area of the corresponding developmental scale before plucking; partial correlation controlling for fish standard length: 0.8, *P* < 10^−10^; dashed line: Total Least Squares, linear fit slope (1.08 ± 0.05), offset (0.18 ± 0.07); n = 131 scales from 19 fish; from 3 experiments. Center and right: scale area at 28 dpp as a function of scale area at 6 dpp (center) and scale area and cell number at 6 dpp (right); 95% CI lower-bound inference for the correlation between cell number at 6 dpp and scale area at 28 dpp: [0.3 0.7]; 95% CI higher bound: [0.96 1]; dashed line: (center: Total Least Squares, linear fit slope (1.57 ± 0.09), offset (−0.1 ± 0.1) mm^2^; right: Ordinary Least Squares with errors in the x-axis, linear fit slope (0.079 ± 0.007) mm^2^, offset (0.18 ± 0.06) mm^2^; center: n = 114 scales from 15 fish; from 2 experiments; right: n = 38 scales from 23 fish from; from 7 experiments (here, new experiments as well as experiments from (*13*) were analyzed)). **D**. Left: number of scale osteoblast nuclei *N* over time in individual scales; right: cell number growth rate (1/*N*) (*dN*/*dt*) (n = 10 scales from 5 fish; from 1 experiment). **E**. Example of fit of cell number over time for an individual scale with a piece-wise linear-constant function (same data as in D as well as experiments from (*13*)) . Root Mean-Square Error (RMSE) is indicated; RMSE = 200 ± 100 in all scales in F. **F**. Relationship between fitted parameters and fitted initial number of cells. Pearson’s correlation coefficient ρ with P-value is indicated; dashed line: Total Least Squares linear fit (center: slope (50 ± 20) h^-1^, offset (-10 ± 40) h^-1^; right: slope (3.2 ± 0.5), offset (0 ± 1); n = 20 scales from 14 fish; from 5 experiments; here, new experiments as well as experiments from (*13*) were analyzed). Scale bar, 200 μm. A-P: anteroposterior axis. Dpp: days post-plucking. Panel A is adapted from (*13*); Copyright © 2021, The Author(s), under exclusive license to Springer Nature Limited.

Scale regeneration allows the study of the coordination of cell and signal dynamics live (*8–13*). Osteoblast-specific promoters (*8, 29–31*) allowed to develop transgenic markers, reporters and biosensors. Using an osteoblast nuclear marker and the fluorescent ubiquitination-based cell-cycle indicator (FUCCI), live microscopy and computational methods, previous work revealed that, at the onset of the proliferative phase, cell division events are distributed roughly uniformly across the bulk of the scale (*9*). Thereafter, cell divisions stop in most of the scale and become restricted to its periphery. However, it is unclear how cell proliferation dynamics and pattern are controlled to obtain a scale with the correct number of cells. In this work, we investigated these questions using a combination of live imaging, transgenic reporters and biosensors, transcriptomics, perturbations and theory.

## Results

We started our analysis by assessing the accuracy of scale size regeneration. To this end, we compared the size of regenerated scales at 28 dpp with the size of the corresponding developmental scales before plucking. We found that, although the size of scales varied across a decade, the size of the regenerated scale strongly correlated with the size of the developmental scale before plucking (Figure 1C left). Furthermore, the number of cells after the proliferative phase (6 dpp), visualized with a transgenic nuclear osteoblast marker, strongly correlated with scale size at the same time, which in turn correlated with regenerated scale size at 28 dpp (Figure 1C center and right). This suggests a tight control of osteoblast number and tissue size in scale regeneration.

To study the control of osteoblast number, we monitored them live. Osteoblast numbers increased roughly linearly over time before reaching a plateau (Figure 1D; 5-6 dpp). We fitted the osteoblast number versus time curves with a piece-wise linear-constant function. We found that the time of transition to plateau did not depend on osteoblast initial number, measured at 2 dpp (Figures 1E and 1F left). Instead, the linear rate of osteoblast number increase scaled with their initial number (Figure 1F center). Altogether, the osteoblast number at plateau (measured at 6 dpp) scaled with their initial number (Figure 1F right). Thereafter, until the completion of regeneration, the number of osteoblasts decreased, likely due to cell death (Figure S1A). Nevertheless, the number of cells just after the end of the proliferative phase (6 dpp) was predictive of regenerated scale size at 28 dpp (Figure 1C). Overall, these results indicate that the initial number of osteoblasts determines their number at the end of the proliferative phase, and thus scale size. In this regenerative process, cell proliferation amplifies osteoblasts by a fixed factor, starting from a scale primordium that is properly scaled to achieve the correct final size. In the simple “counter” and “timer” models, this amplification by a fixed factor can be achieved by maintaining uniform cell proliferation across the tissue and arresting it after a certain number of cell divisions or after a given time. In contrast with these models, the cell number growth rate was not uniform during the proliferative phase, but decreased over time (Fig. 1D), indicating a more complex regulation of cell proliferation.

Given these considerations, we set out to characterize the spatial organization of cell proliferation. To this end, we turned to fish expressing FUCCI in osteoblasts (*9, 32*). FUCCI comprises the G1/G0 phases marker *osx*:zCdt1-mCherry (hereafter, Cdt1) and the S/G2/M phases marker *osx:*Venus-hGeminin (hereafter, Geminin) (Figure 2A left). While entering the cell cycle at the G1/S transition, osteoblasts switch from high levels of Cdt1 and low levels of Geminin (Cdt1^+^ Geminin^−^; G1/G0) to low levels of Cdt1 and high levels of Geminin (Cdt1^−^ Geminin^+^; S/G2/M) (Figures 2A right top (arrowheads), 2B left and S1B; (*9*)). While analyzing FUCCI time courses, we observed a subset of cells in which Cdt1 did not switch off when Geminin turned on, resulting in double-positive Cdt1^+^ Geminin^+^ cells (Figures 2A right bottom (arrowheads), 2B right and S1B). These Cdt1^+^ Geminin^+^ cells did not divide (Figure 2C) and returned to a Cdt1^+^ Geminin^−^ state (Figures 2A right bottom (arrowheads) and 2B right). This behavior is similar to that exhibited by osteoblasts later during regeneration, during the hypertrophic phase, when cells are activated by travelling waves of Erk activity (*13*). Time courses showed that many Geminin^+^ cells were Cdt1^−^ Geminin^+^ early in regeneration (Figure 2D; 2 dpp); thereafter, the fraction of proliferative Cdt1^−^ Geminin^+^ cells decreased progressively, as that of Cdt1^+^ Geminin^+^ cells increased (Figure 2D).

**Figure 2.**
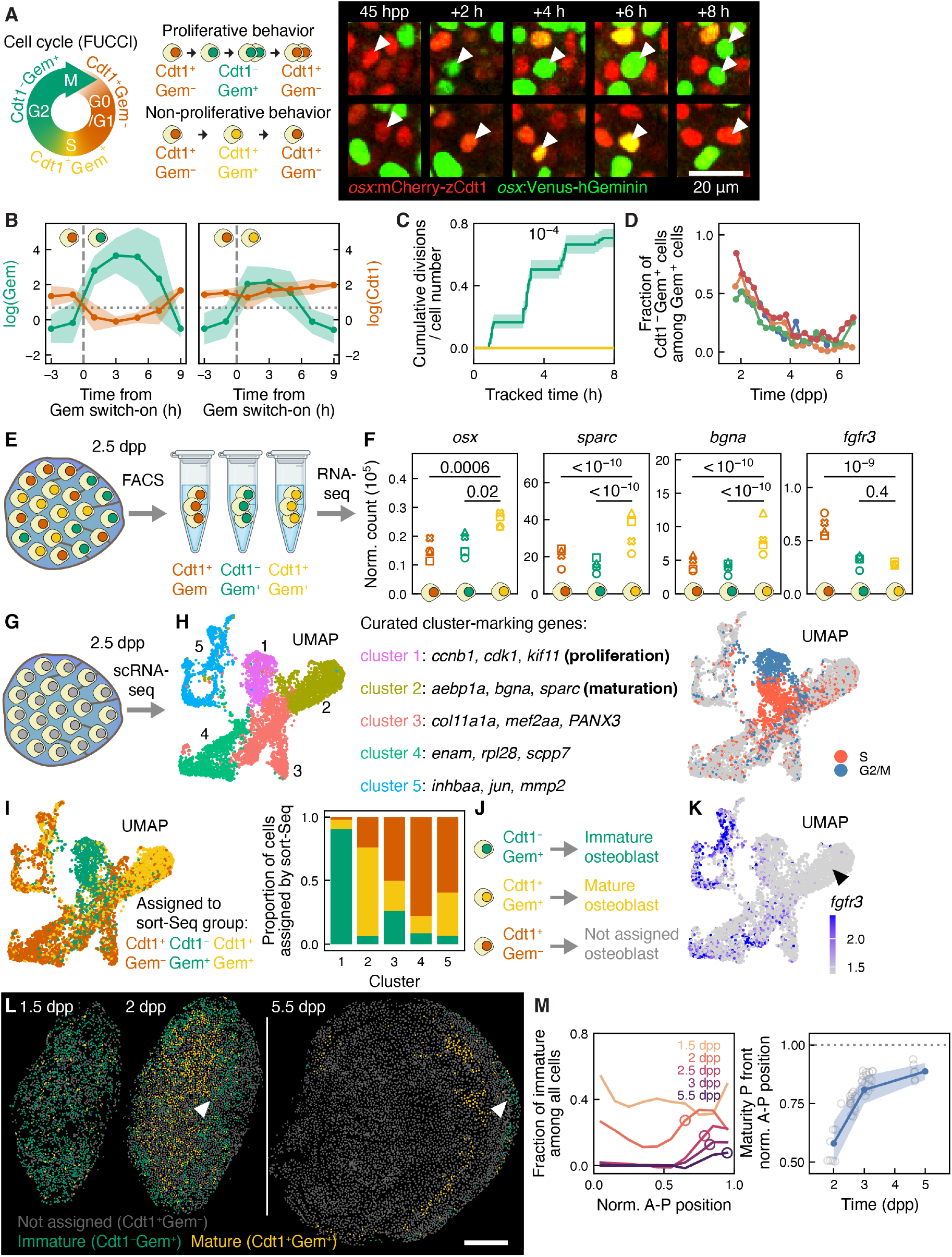
A region of non-proliferative mature osteoblasts expands across the scale. **A**. Schematics of the transgenic cell cycle indicator FUCCI (left), together with schematics (center) and examples (right; arrowhead) of proliferative and non-proliferative behavior (indicated). FUCCI: *osx:mCherry-Cdt1* (hereafter, Cdt1) and *osx:Venus-hGeminin* (hereafter, Geminin). **B**. Cdt1 and Geminin signal in tracked cells exhibiting proliferative behavior (i.e. turning Cdt1^−^ Geminin^+^; left; n = 22 cells from 4 scales from 2 fish; from 2 experiments; with SD across tracks; 1.5–2.5 dpp). Cdt1 and Geminin signal in tracked cells exhibiting non-proliferative behavior (i.e. turning Cdt1^+^ Geminin^+^; right; n = 14 cells from 4 scales from 2 fish; from 2 experiments; with SD across tracks; 1.5–2.5 dpp). **C**. Cumulative number of divisions per tracked cells (Cdt1^−^ Geminin^+^: n = 278 cells; Cdt1^+^ Geminin^+^: n = 12 cells; from 4 scales from 2 fish; from 2 experiments; 2–2.5 dpp). Cumulative division curves were estimated using the Kaplan-Meier estimator (log-rank test P-value is indicated). **D**. Fraction of proliferative osteoblasts (Cdt1^−^ Geminin^+^) over time (n = 4 scales from 4 fish; from 1 experiment; re-analyzed from (*13*)). **E**. Schematics of bulk mRNA sequencing (RNAseq) experiment following sorting (sortSeq). FACS: fluorescence-activated cell sorting. **F**. Normalized transcript counts (DESeq2) in sortSeq experiment for select osteoblast genes (40–60 scales at 2.5 dpp from 2-3 fish per sample; from 1 experiment). DESeq2 paired adjusted P-value is indicated. **G**. Schematics of single-cell mRNA sequencing (scSeq) of sorted *osx:*H2A-mEos2^+^ osteoblasts (scSeq; 40–60 scales at 2.5 dpp from 9 fish; from 1 experiment). **H**. Left: scSeq clusters. UMAP was calculated on the first 20 principal components of the data which capture 80% of the total variance. Right: cell cycle score (Methods). **I**. Visualization (left) and quantification (right) of assignment of scSeq cells to sortSeq cell pools based on detected transcripts in each experiment. **J**. Schematics of assignment of osteoblast populations detected with FUCCI to immature and mature populations detected with transcriptomics. **K**. Visualization of *fgfr3* transcripts per cell in scSeq experiment. Color bar: Seurat log-normalized transcript counts. Arrowhead: maturation cluster. **L**. Example of distribution of immature Cdt1^−^ Geminin^+^ and mature Cdt1^+^ Geminin^+^ osteoblasts over time. Arrowhead: maturation front. **M**. Left: example of profile of fraction of immature Cdt1^−^ Geminin^+^ osteoblasts across the A-P axis over time (same scales as in L; circles: front position). Right: position of the posterior front of the mature region over time (circles: front position for each scale and time-point from n = 27 scales from 13 fish; from 7 experiments; with SD; hereafter, front position is scored only when detectable, see Figure S2A). Scale bars, 200 μm, unless differently indicated. A-P: anteroposterior axis. Dpp: days post-plucking. Gem: Geminin. Norm: normalized. P front: posterior front.

Since Cdt1^+^ Geminin^+^ osteoblasts showed a similar behavior as osteoblasts during the hypertrophy phase, we hypothesized that non-proliferative Cdt1^+^ Geminin^+^ osteoblasts were terminally differentiated mature cells, compared with proliferative Cdt1^−^ Geminin^+^ osteoblasts, who would be immature. To investigate these hypotheses, we analyzed the osteoblasts’ transcriptome following fluorescence-activated cell sorting (FACS) (Figure 2E; Methods). We found that, compared with Cdt1^−^ Geminin^+^ and Cdt1^+^ Geminin^−^ cells, Cdt1^+^ Geminin^+^ osteoblasts exhibited higher transcript levels of the osteoblast maturation transcription factor *osterix*, as well as the bone formation genes *biglycan a (bgna)* and *osteonectin (sparc)* (Figure 2F). Conversely, compared with Cdt1^+^ Geminin^−^ cells, Cdt1^+^ Geminin^+^ osteoblasts, showed lower transcripts levels of the *fibroblast growth factor receptor 3* (*fgfr3)* (Figure 2F), whose protein hyperactivation impairs bone development in achondroplasia (*33, 34*).

To further investigate the hypothesis that the Cdt1^−^ Geminin^+^ and Cdt1^+^ Geminin^+^ cells were cells at different states of maturation, we analyzed the transcriptome of individual osteoblasts using single-cell RNA sequencing (Figures 2G, 2H and S1C; 2.5 dpp; scRNA-seq; Methods). After clustering individual cells based on their transcriptome, we identified five clusters (Figure 2H). One such cluster expressed S and G2/M markers (Figure 2H left, cluster 1 and Figure 2H right). Instead, another one expressed markers of osteoblast maturation (Figure 2H left, cluster 2). To test whether these two clusters corresponded to the Cdt1^+^ Geminin^+^ and Cdt1^−^ Geminin^+^ populations identified using the FUCCI system, we mapped the transcriptome of individual cells to the bulk transcriptome of the sorted proliferative and non-proliferative populations we previously obtained. Strikingly, the cluster of mature osteoblasts mapped mainly to the non-proliferative Cdt1^+^ Geminin^+^ population, while the cluster of proliferative osteoblasts mapped mainly to the proliferative Cdt1^−^ Geminin^+^ population (Figures 2I and 2J). Cdt1^+^ Geminin^−^ cells were the most represented in the remaining three clusters. As we found in the case of bulk RNAseq, mature osteoblasts expressed low levels of *fgfr3* (Figure 2K (arrowhead)). We concluded that Cdt1^+^ Geminin^+^ osteoblasts were more mature than Cdt1^−^ Geminin^+^ osteoblasts.

Following up on these observations, we set out to investigate the appearance of mature Cdt1^+^ Geminin^+^ osteoblasts over time (Figures 2L and 2M). We discovered that, initially, a region enriched in mature cells formed centrally in the scale (Figures 2L center, 2M, S2A and S1B). Thereafter, this mature region expanded across the scale, progressively restricting most immature Cdt1^−^ Geminin^+^ cells to the scale boundary (Figures 2L right, 2M, S2A and S1B). Thus, a moving front of osteoblast maturation traversed the regenerating scale at the transition from proliferation to hypertrophy. In this context, the front is defined as the transition point from a region of immature cells to a region of mature ones (Methods). As expected, the immature region showed higher occurrence of cell divisions, while the mature region showed a higher rate of cell growth (Figures 3A and 3B). Overall, these results revealed that a mature, non-proliferative and hypertrophic region emerges from the scale center and progressively expands throughout the scale. This mature region can be detected using FUCCI for having high occurrence of Cdt1^+^ Geminin^+^ cells.

**Figure 3.**
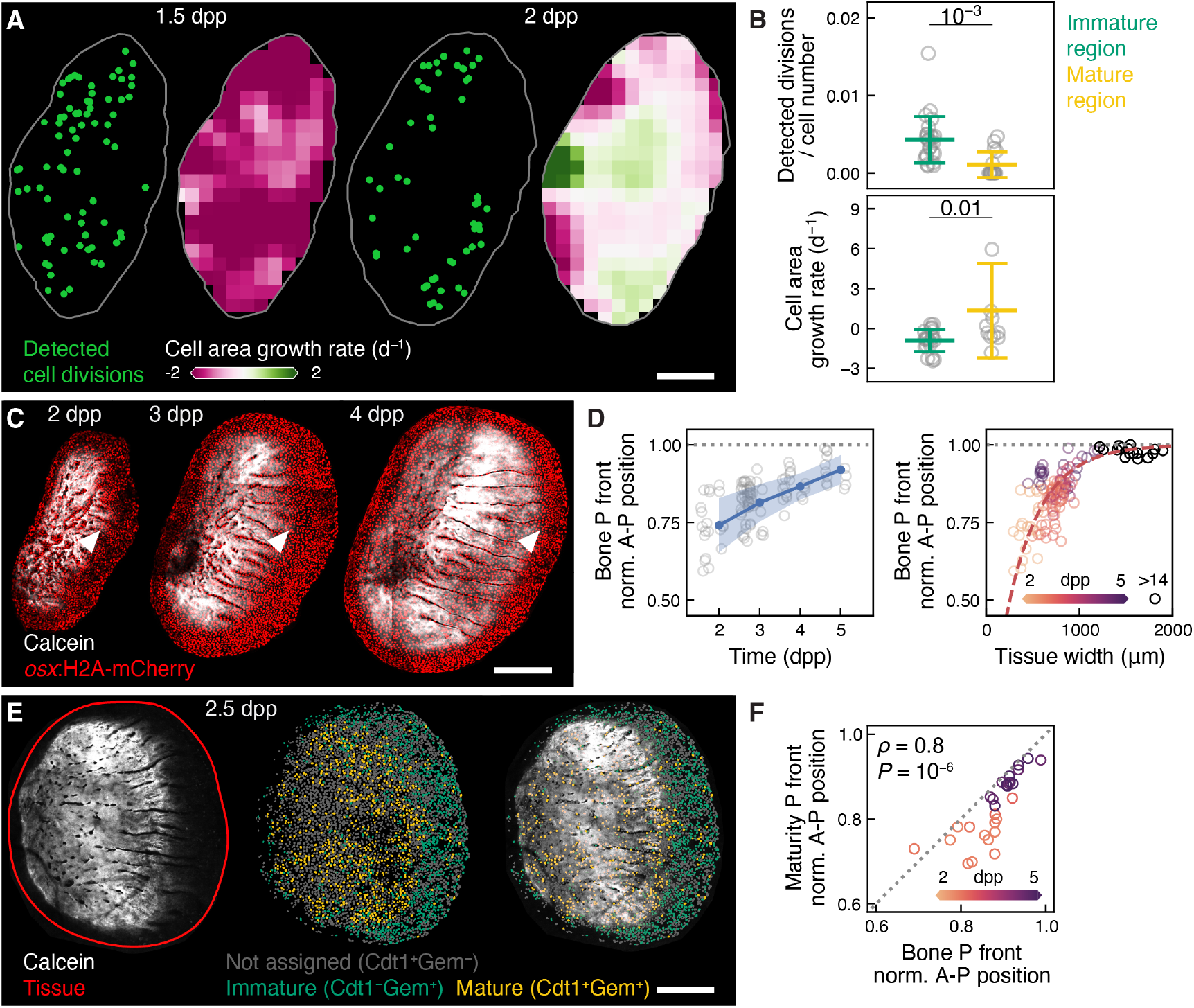
The expanding mature osteoblast region is associated with hypertrophic behavior and bone deposition. **A**. Example of distribution of detected cell divisions and cell area growth rate map over time (same scale as in Figures 2L and 2M left). **B**. Detected cell divisions per cell (top) and cell area growth rate (bottom) in the immature and mature scale regions (circles: scale-wide averages of time points from n = 4 scales from 2 fish; from 2 experiments; with SD). Student’s unpaired t-tests’ P-values are indicated. **C**. Example of Calcein bone staining (white) with osteoblast nuclei (red) over time. Arrowheads: bone front. **D**. Position of the posterior front of the bone (Calcein or Alizarin staining) over time (left; circles: individual scales and time points from n = 66 scales from 28 fish; from 11 experiments; with SD) and over scale A-P tissue width (right; circles: time points from n = 90 scales from n = 35 fish; from 14 experiments; dashed line: saturating exponential fit). **E**. Example of Calcein bone staining and osteoblast maturity. Solid red line: scale contour. **F**. Position of the posterior front of the mature region as a function of the position of the posterior front of the bone (Calcein or Alizarin staining) (circles: individual scales; n = 28 scales from n = 12 fish; from 5 experiments; Pearson’s correlation coefficient ρ with P-value is indicated; dashed line: bisector of the axes). Scale bars, 200 μm. A-P: anteroposterior axis. Dpp: days post-plucking. Gem: Geminin. Norm: normalized. P front: posterior front.

Then, we tested if the appearance of the mature region was also associated with bone deposition, as typical for mature osteoblasts (*16, 20*). Thus, we compared the extent of the mature region with that of the mineralized scale bone stained with the live dye Calcein (hereafter: the bone). At 2 dpp, the bone region already covered a significant portion of the scale; thereafter, the bone region expanded until covering most of the scale at 5 dpp (Figures 3C and 3D). Strikingly, we found that the bone deposition front (Methods) correlated with the front of establishment of the mature region, enriched in Cdt1^+^ Geminin^+^ cells (Figures 3E and 3F). Thus, osteoblast maturation was associated with mineralized bone deposition. Intriguingly, we observed that, at early time points, the bone deposition front was slightly advanced with respect to the maturation front (Figure 3F). This raises the possibility that early bone, deposited by immature osteoblasts, may contribute to osteoblast maturation.

Thereafter, we interrogated what signaling dynamics were associated with osteoblast maturation. In scales, Fibroblast Growth Factor (Fgf) signaling is necessary for osteoblast proliferation and was suggested to be involved in the patterning of cell proliferation (*9*). Compatible with this notion, *fgf20a* expression switches from a uniform pattern to being restricted to the scale boundary (*9*). Furthermore, activation of the Fgfr downstream kinase Erk showed a similar transition (*13*), as revealed by the Erk activity biosensor *osx:*ErkKTR-mCerulean (hereafter: ErkKTR) (*13, 35*). Therefore, we characterized the dynamics of Erk activity as it transitioned to being restricted to the periphery. To this end, we performed high time-resolution imaging of scales expressing the ErkKTR biosensor. These experiments revealed a moving front of Erk inactivation that progressively travelled across the tissue (Figures 4A, 4B and S2A; Methods). We noted that Erk activity slightly decreased in the Erk-active region over time, indicating an Erk temporal dynamics additional to the moving front of inactivation (Figure 4C). We compared the pattern of osteoblast maturation with that of Erk inactivation and we found that the two fronts coincided (Figure S3A). Accordingly, an analysis of the cell division spatial distribution showed fewer detected cell divisions in the low Erk activity region compared with the high Erk activity one (Figure 4A top and 4D). Finally, pharmacological inhibition of Erk activity strongly reduced osteoblast number growth rate (Figures S3B, S3C, S3D and S3E top). In addition, this Erk inhibition reduced Cdt1^+^ Geminin^+^ across the scale (Figures S3D and S3E), indicating that Erk activity is necessary for cells to temporarily turn from Cdt1^+^ Geminin^-^to Cdt1^+^ Geminin^+^. Altogether, these observations indicate that Erk activity is required for osteoblast proliferation and that, starting centrally, a front of Erk inactivation and arrest of proliferation propagates across the scale. However, thereafter, Erk is activated in travelling waves that do not trigger cell proliferation (Figures S4A and S4B) (*13*). Thus, Erk activation is required, but not sufficient, for cell proliferation.

**Figure 4.**
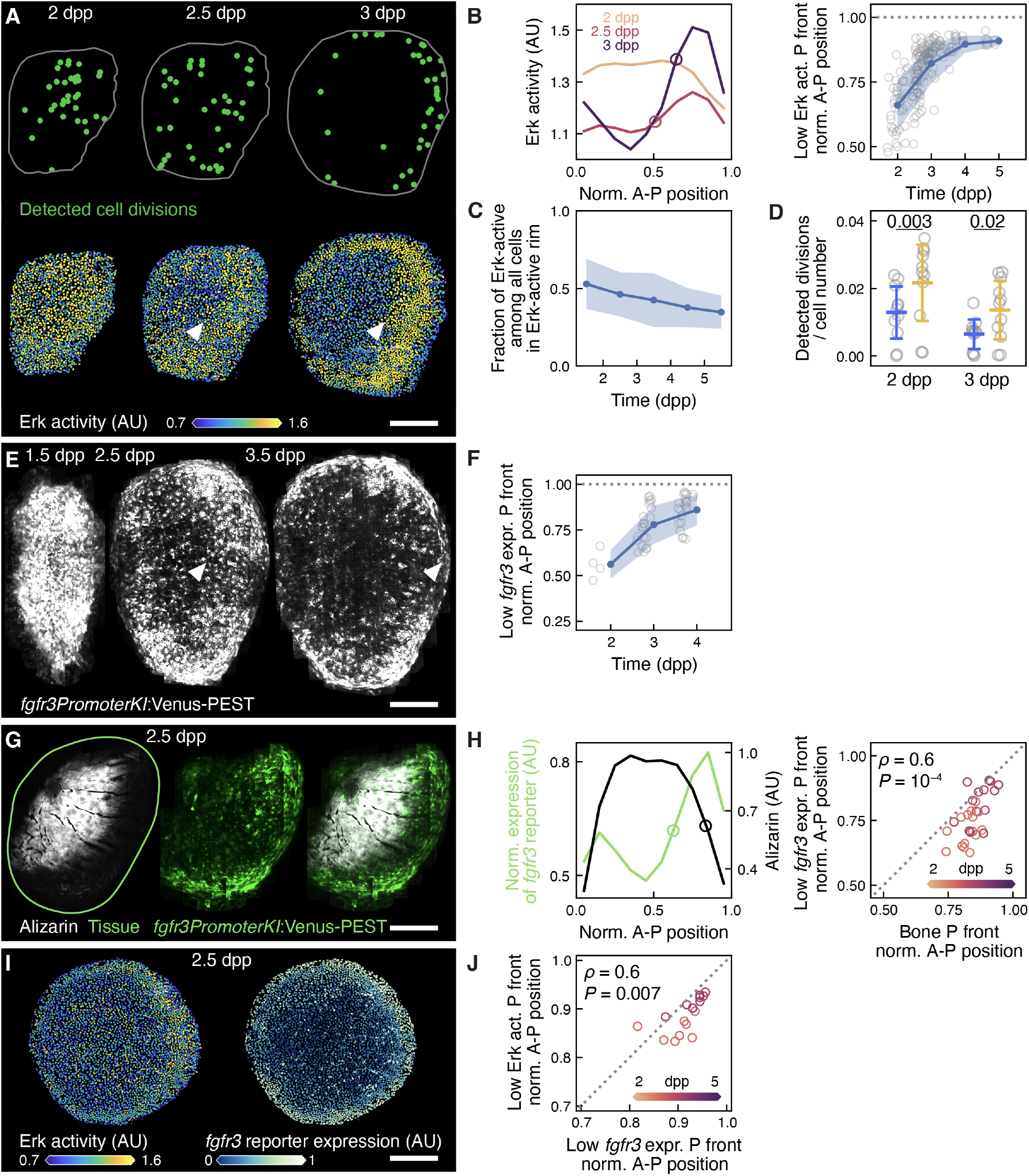
A moving front of Erk inactivation and reduction of *fgfr3* expression accompanies osteoblast maturation. **A**. Example of detected cell divisions (top) and Erk activity (bottom). Arrowheads: Erk inactivation front. **B**. Left: example of Erk activity profile across the A-P axis in an individual scale over time (same scale as in A; circles: front position). Right: quantification of the position of the posterior front of the low Erk activity region over time (circles: individual scales and time points from n = 77 scales from 38 fish; from 19 experiments; with SD). **C**. Fraction of Erk-active cells in the immature region over time; linear fit with slope (−0.06 ± 0.01) d^−1^ (n = 83 scales from 43 fish; from 20 experiments; with SD). **D**. Detected cell division number per cell in low and high Erk activity regions (circles: individual scales; n = 11 scales from 6 fish; from 4 experiments; with SD). Student’s paired t-tests’ P-values are indicated. **E**. Example of signal of the *fgfr3PromoterKI*:Venus-PEST transcriptional reporter (hereafter, *fgfr3* reporter). Arrowheads: front. **F**. Position of the posterior front of the of the region of low *fgfr3* expression over time (circles: individual scales and time points from n = 24 scales from 9 fish; from 3 experiments; with SD). **G**. Example of *fgrf3* reporter expression and Alizarin bone staining. **H**. Left: profile of *fgfr3* reporter expression and bone (Alizarin staining) staining across the A-P axis over time (same scale as in G; circles: front position). Right: position of the posterior front of the of the region of low *fgfr3* expression as a function of the position of the posterior front of the bone region (circles: individual scales and time points from n = 16 scales from 5 fish; from 2 experiments; Pearson’s correlation coefficient ρ with P-value is indicated; dashed line: bisector of the axis). **I**. Example of Erk activity and quantified *fgfr3* reporter expression. **J**. Position of the posterior front of the low Erk activity region as a function of the position of the posterior front of quantified low *fgfr3* expression (circles: individual scales and time points from n = 8 scales from 4 fish; from 1 experiment; Pearson’s correlation coefficient ρ with P-value is indicated; dashed line: bisector of the axis). Scale bars, 200 μm. A-P: anteroposterior axis. AU: arbitrary units. Dpp: days post-plucking. Expr.: expression. Gem: Geminin. Norm: normalized. P front: posterior front.

Since our transcriptome analysis identified *fgfr3* expression as a marker of osteoblast maturation, a reduction of *fgfr3* expression could in principle explain the reduction of Erk activity in those osteoblasts. Thus, we decided to monitor *fgfr3* expression throughout the scale over time. To this end, we generated a new *fgfr3* transcriptional reporter *(fgfr3KI*:Venus-PEST) using an endogenous promoter Crispr Knock-In technique (*36, 37*). Time courses of this newly established reporter showed that *fgfr3* was expressed throughout the scale at the onset of the proliferative phase (Figures 4E and 4F). Thereafter, *fgfr3* expression turned off in a propagating front (Figures 4E, 4F and S2A; Methods). The expanding region of *fgfr3* inactivation matched with the maturation region, detected by bone deposition (Figures 4G and 4H; Alizarin) and Erk inactivation (Figures 4I and 4J).

In addition to Erk, the p38 kinase is an important signaling hub involved in osteoblast proliferation and maturation (*26*). Therefore, we asked if p38 activity pattern was related to the progressive maturation of the scale osteoblast tissue. To monitor p38 activity, we implemented a p38 KTR biosensor (*35*) in zebrafish (*osx*:p38KTR-Venus; Figure 5A left). To validate this newly generated sensor line, we performed immunostainings using an antibody for the active, i.e. phosphorylated, form of p38 (anti-p-p38; Figure 5A right). Comparing the p38KTR readout with anti-p-p38 staining revealed a linear sensor response covering most of the p38KTR dynamic range (Figure 5B). Intriguingly, live imaging showed that p38 inactivated in a propagating front that expanded throughout the tissue (Figures 5C, 5D and S2A; Methods). The fronts of p38 and Erk inactivation matched (Figure 5C and 5E). Overall, these findings showed that osteoblast maturation is accompanied by Erk and p38 inactivation.

**Figure 5.**
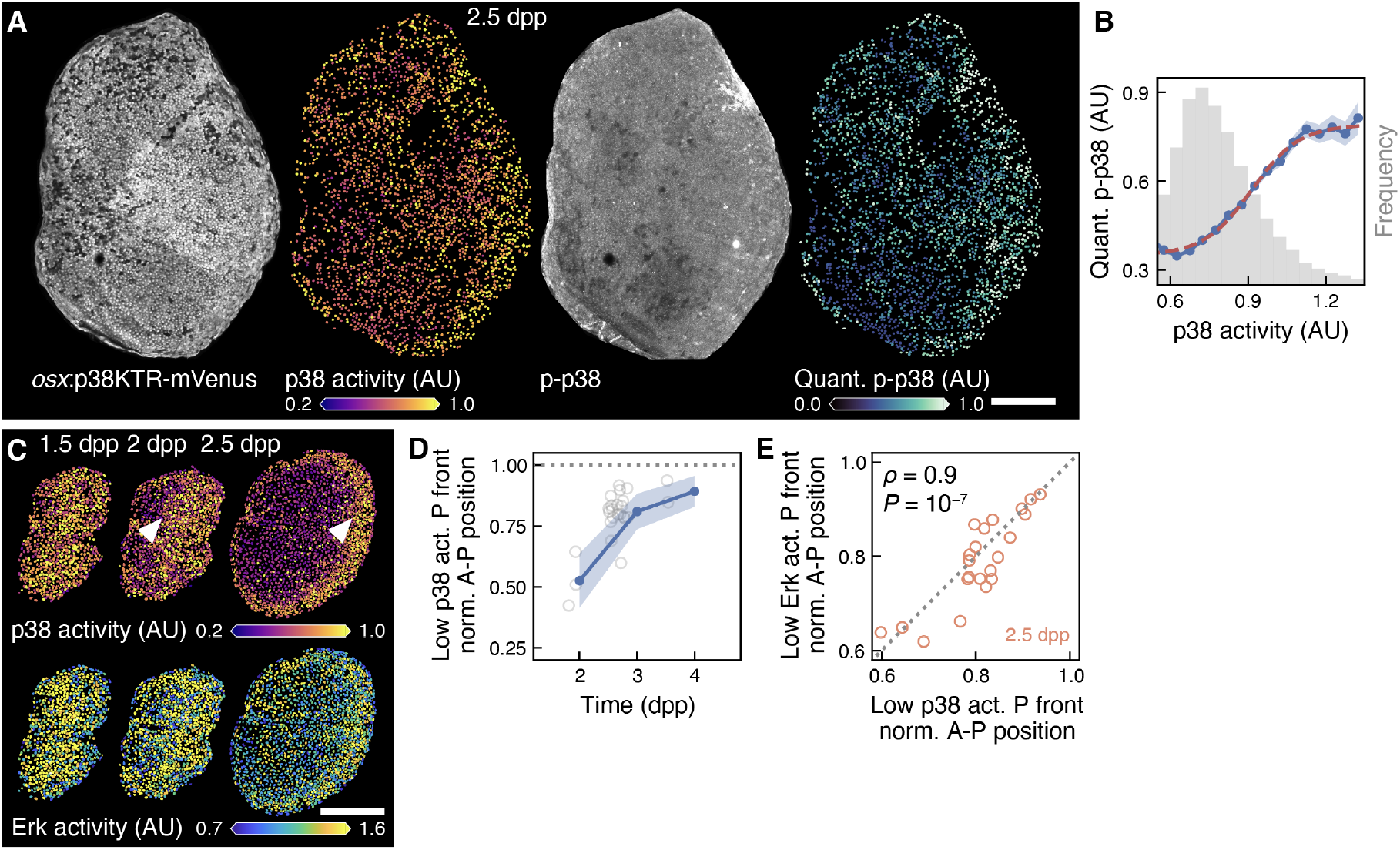
A new p38 activity biosensor fish reveals a moving front of p38 inactivation that accompanies osteoblast maturation. **A**. Example of p38KTR signal, quantified p38 activity, anti-p-p38 immunostaining and quantified anti-p-p38. **B**. Quantified anti-p-p38 immunostaining as a function of p38 activity reported by the p38KTR sensor (n = 7 scales from 3 fish; from 1 experiment; with SEM; dashed line: sigmoidal fit; grey: frequency of observed p38 activity values). **C**. Example of p38 and Erk activity over time. Arrowheads: p38 inactivation front. **D**. Position of the posterior front of the low p38 activity region over time (circles: individual scales and times points from n = 18 scales from 9 fish; from 4 experiments; with SD). **E**. Position of the posterior front of the low p38 activity region as a function of position of the posterior front of the low Erk activity region (circles: individual scales and time points from n = 18 scales from 9 fish; from 4 experiments; Pearson’s correlation coefficient ρ with P-value is indicated; dashed line: bisector of the axis). Scale bars, 200 μm. A-P: anteroposterior axis. AU: arbitrary units. Dpp: days post-plucking. Norm: normalized. Quant: quantified. P front: posterior front.

Our results show that osteoblast maturation is associated with proliferation arrest. Thus, we hypothesized that maintaining osteoblasts in an immature state would enhance their proliferation. Serendipitously, we observed that maintaining osteoblast immaturity could be achieved by temporally over-expressing Fgf20a using a heat-shock inducible promoter during the proliferative phase (Figures 6A and 6B). This perturbation caused some osteoblasts to migrate away from the scale during the 24 h following induction (Figure S4C), potentially indicating a return to a mesenchymal state. Thereafter, two days after induction, among the osteoblasts that stayed in the scale, the overall fraction of Erk^+^ and Geminin^+^ cells returned to similar levels as in control scales (Figures 6A and 6B top). Nevertheless, we detected a higher frequency of immature Cdt1^−^ Geminin^+^ cells compared with control (Figures 6A and 6B bottom left). Strikingly, such elevation in immature cells was associated with higher osteoblast number increase (Figure 6B bottom right). Heat-shock performed later in regeneration, during the hypertrophic phase, could activate Erk immediately thereafter, but the fraction of immature osteoblasts and osteoblast number increase was not elevated, neither immediately after induction (Figures S4D and S4E) nor two days after (Figures S4F and S4G). These results support the model in which osteoblast maturation halts their proliferation, which cannot be re-activated by Erk during the hypertrophic phase.

**Figure 6.**
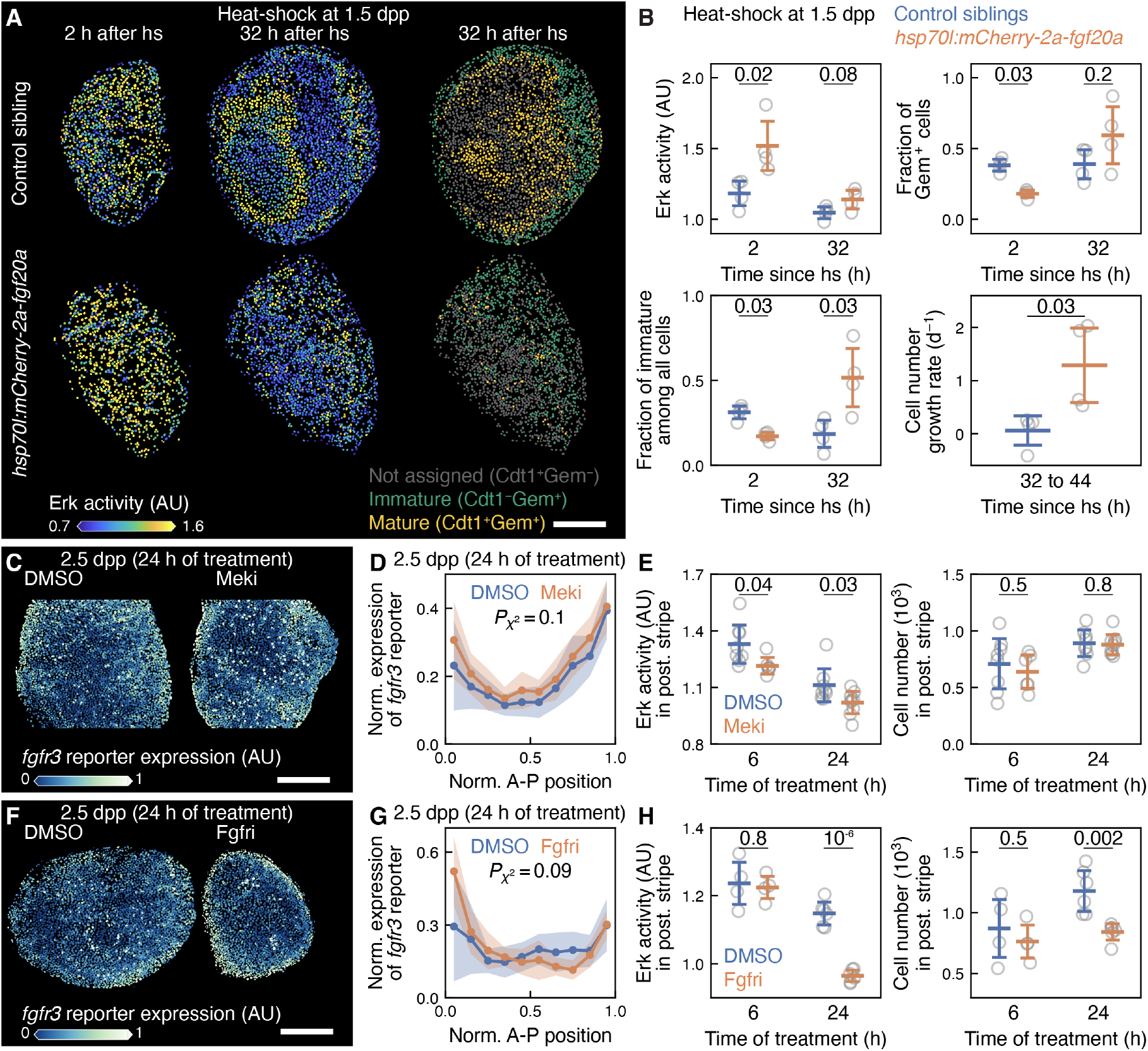
Experimental perturbations maintaining osteoblasts in an immature state enhance cell proliferation. The moving front is not impaired by inhibition of Fgfr/Erk signaling and cell proliferation. **A**. Example of Erk activity and mature/immature cells following heat-shock at 1.5 dpp in *hsp70l:mCherry-2a-fgf20a* and control fish. **B**. Erk activity, fraction of Geminin*^+^*cells, fraction of immature cells and cell number growth rate following heat-shock at 1.5 dpp in *hsp70l:mCherry-2a-fgf20a* and control fish (circles: individual scales; control n = 4 scales from 2 fish, *hsp* n = 4 scales from 2 fish; from 1 experiment; with SD; Student’s unpaired t-tests’ (Erk activity and proliferation) and Mann-Whitney U tests’ (fractions) P-values are indicated). **C**, **D**, **E**. Example (C) and profile quantification across the A-P axis of *fgfr3* reporter expression (D, with SD across scales), and Erk activity and cell number in posterior stripe in control and treated (Mek inhibitor PD0325901, 10 μM) fish (E; circles: individual scales; with SD; DMSO n = 8 scales from 4 fish, Meki n = 6–9 scales from 3 fish; from 1 experiment). **F**, **G**, **H**. Example (F) and profile quantification across the A-P axis of *fgfr3* reporter expression (G, with SD across scales), Erk activity and cell number in posterior stripe in control and treated (pan-Fgfr inhibitor BGJ398, 10 μM) fish (H; circles: individual scales; with SD; DMSO n = 4–6 scales from 2–3 fish, Fgfri n = 4–6 scales from 2–3 fish ; from 1 experiment). χ2 tests’ (D, G) and Student’s unpaired t-tests’ (E, H) P-values are indicated. Scale bars, 200 μm. A-P: anteroposterior axis. AU: arbitrary units. Dpp: days post-plucking. Gem: Geminin. Norm: normalized. Post: posterior. P front: posterior front.

Since osteoblast maturation was associated with reduction of Fgfr/Erk signaling, we asked if loss of Fgfr/Erk signaling itself would perturb osteoblast maturation. We found that pharmacological Fgfr/Erk inhibition did not significantly impair the osteoblast maturation pattern, measured using the pattern of *fgfr3* expression (Figures 6C–H). Interestingly, we noted that osteoblast maturation was not perturbed in spite of the impairment of osteoblast proliferation resulting from Fgfr inhibition (Figures 6F–H). Thus, osteoblast maturation proceeded regardless of inhibition of Fgfr/Erk signaling and proliferation.

Our results show that the moving front of osteoblast maturation spatially restricts and thus progressively halts cell proliferation, traversing each portion of the scale at a given regeneration time (Figure 7A–C and S4H). Intriguingly, the dynamics of osteoblast maturation scales with tissue size, to reach the tissue end at the same time in all scales (Figures 7B, S4H and S4I).

**Figure 7.**
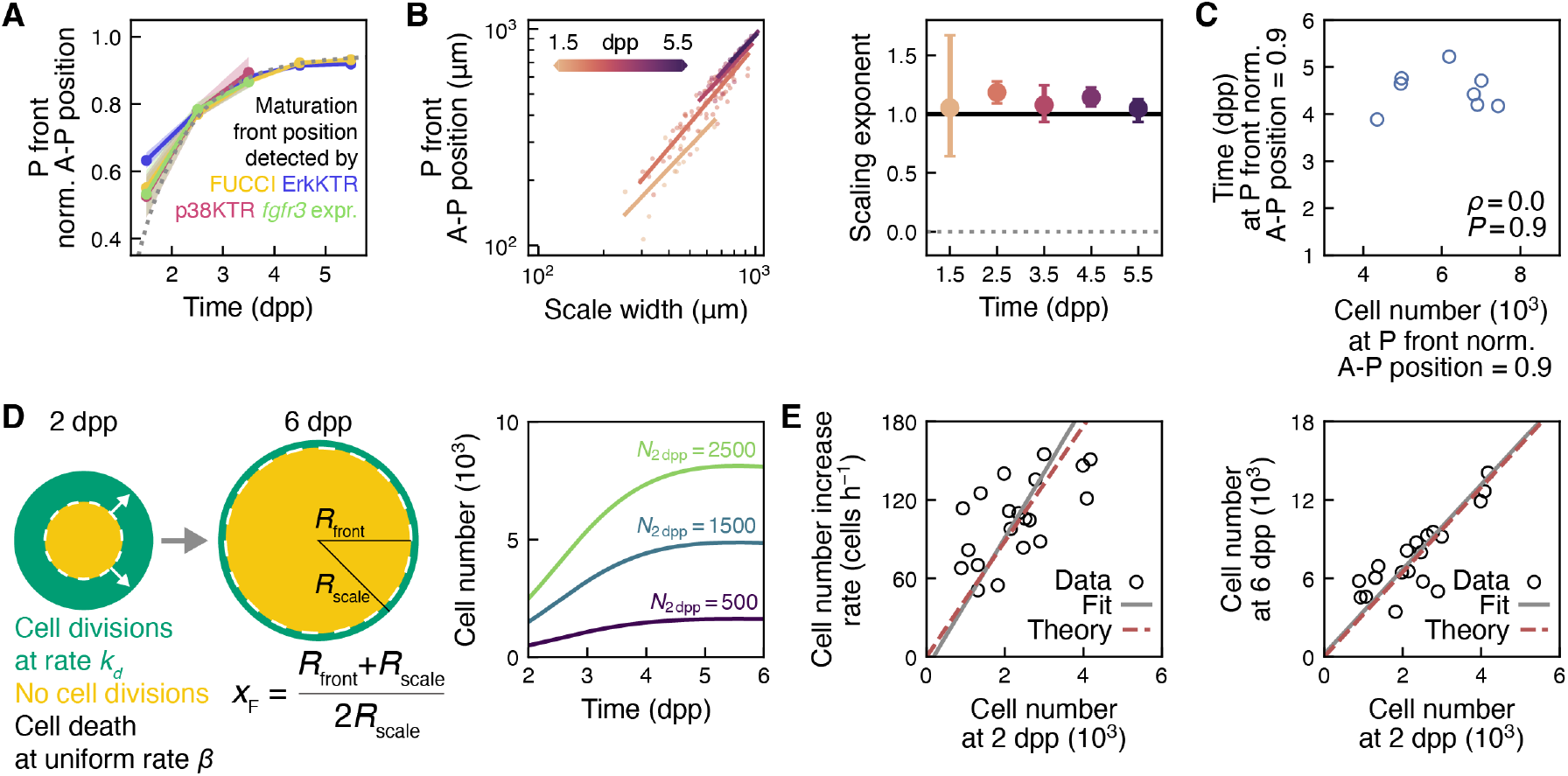
The osteoblast maturation front scales proliferation dynamics and final osteoblast number. **A**. Overview of maturation front x_F_ dynamics obtain with different measurements of maturation (exponential relaxation fit x_F_ = x_F, MAX_ + (x_F,0_ − x_F, MAX_)·exp(−(t − 48h)/τ), x_F,0_ = 0.639 ± 0.008, x_F,max_ = 0.95 ± 0.01, and τ = (24 ± 2) h; with SEM; data from Figures 2M, 4B, 4F and 5D, as well as experiments from (*13*) for ErkKTR and FUCCI). **B**. Left: Position of the posterior front of maturation as a function of scale width (from A; with power-law fits). Right: slope of fits from center right with 95% confidence interval. **C**. Time at which the maturation front has traversed 90% of the scale compared with cell number at that time (circles: individual scales; Pearson’s correlation coefficient ρ with P-value is indicated; from A). **D**. Mathematical model of osteoblast number dynamics. Left: schematics. Right: cell number as a function of time in the mathematical model of osteoblast proliferation and death dynamics (N_2 dpp_ is the number of cells at the initial condition). **E**. Cell number increase rate *dN*/*dt* and cell number at 6 dpp as a function of cell number at 2 dpp in the mathematical model of proliferation and death dynamics (with data and fits from Figure 1F).

We wondered whether such scaling dynamics of the moving front could explain the osteoblast proliferative dynamics and its scaling with osteoblast number that we had initially observed (Figure 1F). To test this, we implemented a mathematical model that assumed that the front separated a proliferating region with a uniform constant rate of proliferation from a non-proliferating region and included a basal rate of cell death (Figure 7D left). We found that this model, together with the observed front dynamics, could explain the osteoblast number dynamics, as well as the scaling of its slope and plateau with osteoblast initial number (Figures 7D right and 7E; Supplementary Information).

## Discussion

In summary, live imaging, transcriptomics, quantifications and theory reveal a moving front of osteoblast maturation that progressively traverses the scale tissue. Maturation, as it travels across the scale, switches osteoblasts from a proliferative to a hypertrophic behavior. The maturation of osteoblasts is accompanied by decrease in p38 activity, *fgfr3* expression and, accordingly, of baseline Erk activity. However, the experimental reduction of Fgfr/Erk signaling could not accelerate osteoblast maturation itself.

Our findings show that the scale original size is recovered by controlling two components: the size of the initial tissue, which scales with the scale original size, and the factor of amplification of the initial osteoblast pool. The size of the initial tissue could be set appropriately by the size of scale pocket from where the scale was plucked and in which regeneration takes place. Complementarily, we find that osteoblast amplification is controlled by the dynamics of the moving maturation front, which is the same in all scales, once scaled with tissue size. Remarkably, the front position continued to scale even after impairing Erk activity and thus cell proliferation. As a result, the maturation front causes osteoblast number to reach a plateau at the same time. However, differently from the simple “timer” model, osteoblasts stop proliferating at different times across the scale as the front progressively travels. In contrast, we propose a “scaler” model, in which a maturation front dynamics that scales with tissue size scales cell number by a fixed factor. Similar proliferative control by osteoblast maturation may take place in the zebrafish fin, in which cell proliferation and Fgf/Erk signaling decrease and become progressively restricted to the tissue proliferative edge as regeneration proceeds (*38*).

The dynamics of the moving front raises questions regarding how it is initiated and propagated across the scale in a way that is the same across scales, once scaled with tissue size. A moving front restricting cell proliferation has been described in development in the larval *Drosophila* eye imaginal disk (*39–41*). In this system, a differentiation front, located at a moving tissue structure called “morphogenetic furrow”, propagates from the posterior to the anterior of the tissue and meanwhile arrests cell proliferation. Morphogen gradients move together with the differentiation front and modulate cell proliferation in the entire tissue (*42, 43*). Such propagation is mediated partly by Hedgehog signaling and behaves as a chemical trigger wave (*44–47*). Differently from the front observed in scales, chemical trigger waves move at a constant speed and thus their dynamics would not scale with tissue size. In the fly eye disk, a second mechanism limits cell proliferation, since the rate of cell proliferation decreases in the undifferentiated and proliferating region of the disk (*48*). This reduction has been proposed to be driven by the progressive dilution of the chemokine Upd due to tissue growth (*49*), thus providing an effective sizer mechanism (*48*). A similar mechanism could be active in the scale to adapt the dynamics of the moving front to tissue size.

Similarly to the classic “mechanical contact inhibition” model (*5, 6, 50–53*), osteoblast maturation could be triggered by forces rising as cells become denser. In this case, cell density could in principle provide information for scaling the front position with tissue size. Alternatively, the stiffness of the bone matrix could provide information regarding the progression of regeneration and contribute to determining front position. Indeed, we found that the maturation of osteoblasts is associated with the deposition of calcified bone matrix. This result indicates that the bone itself may play a role in osteoblast maturation, as recently observed in mouse calvarial osteoblasts (*54*). The development of targeted bone matrix and mechanical perturbations will be needed to distinguish between these models *in vivo*.

Altogether, this work showed a moving front of osteoblast maturation, associated with reduction of p38 and Fgf/Erk signaling, which controls proliferation during zebrafish scale regeneration. These events are highly stereotypically coordinated across the osteoblast monolayer in different scales and animals. Similar maturation-based mechanisms may occur in other systems in which cellular events need to be rapidly coordinated across large regenerating tissues.

## METHODS

### Key Resources Table

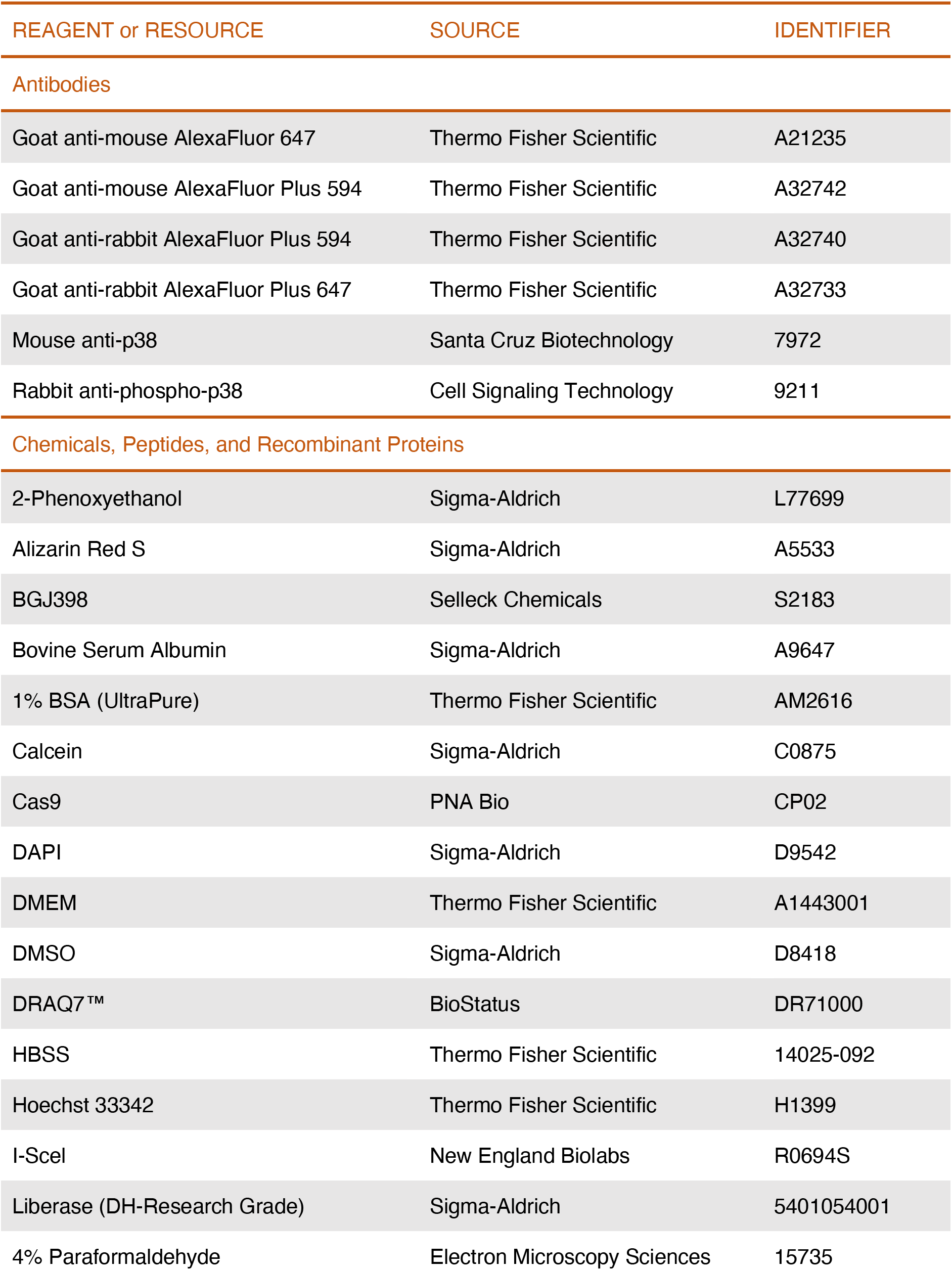
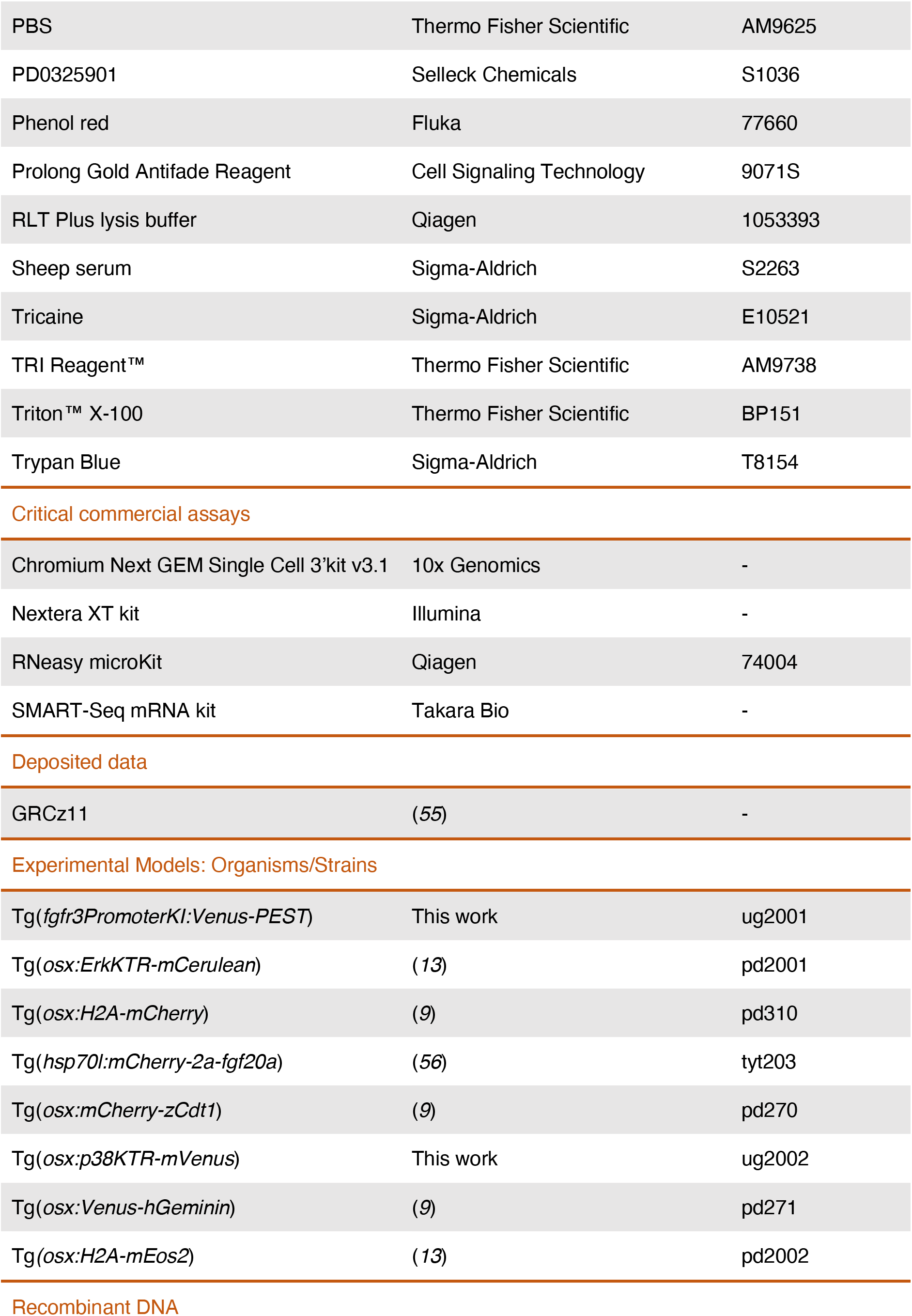
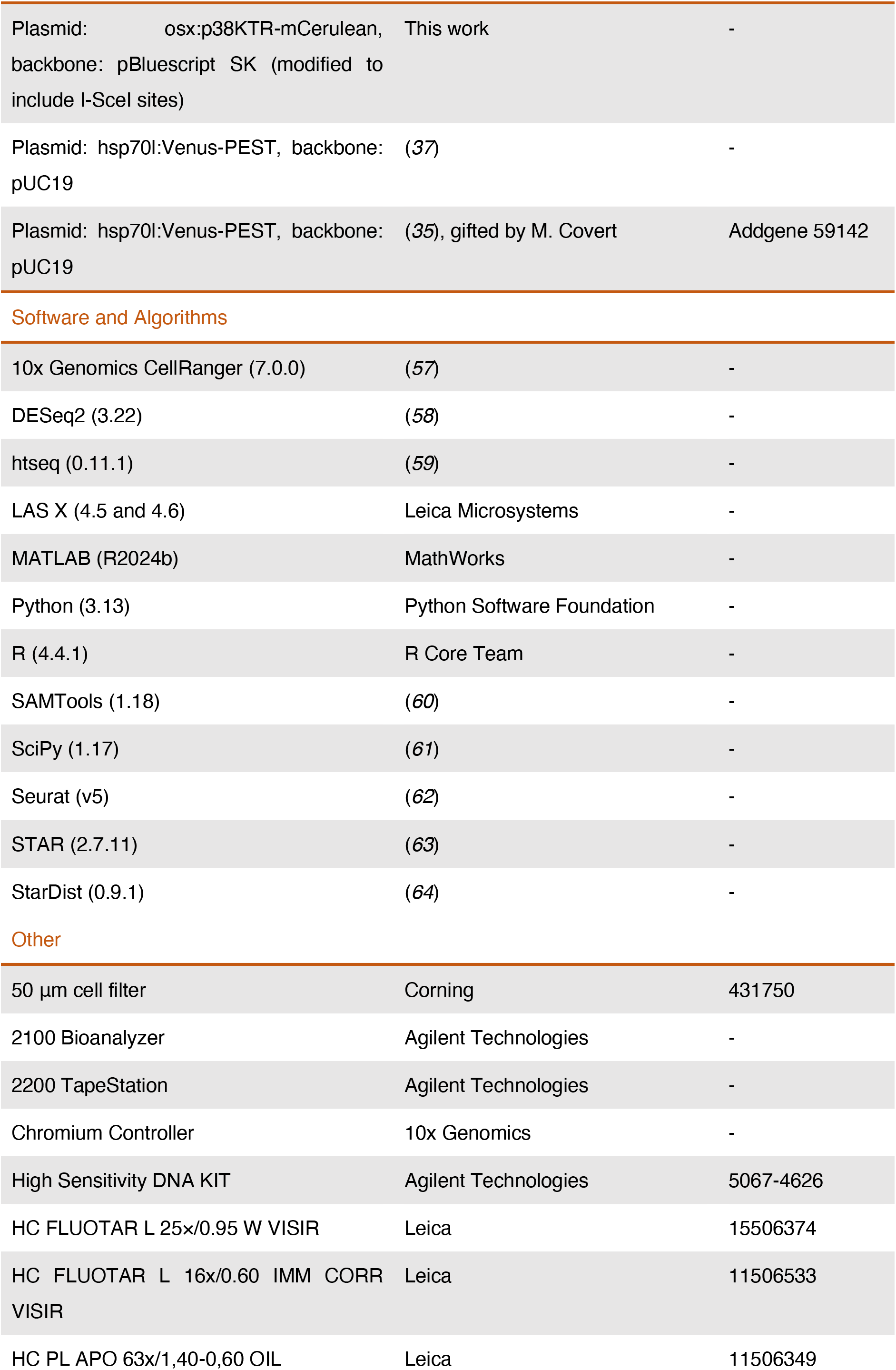
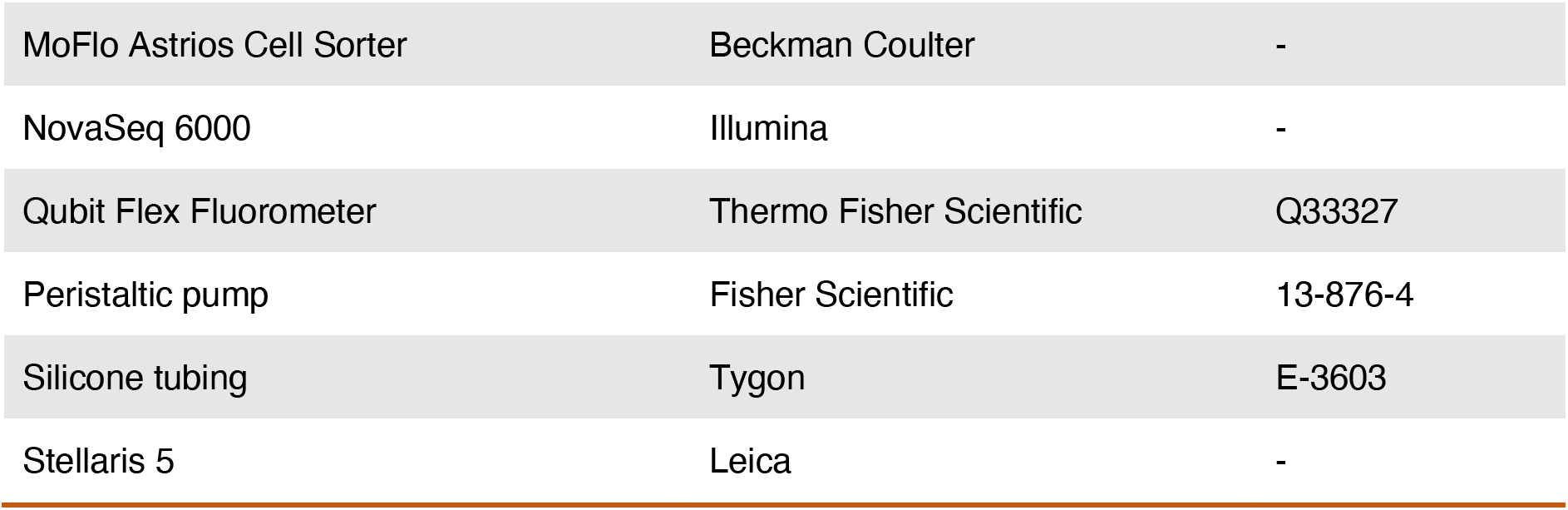

### Fish husbandry, scale injury and pharmacological treatments

Zebrafish of Ekkwill, Ekkwill/AB and AB* strains were maintained between 26 and 28.5 °C with a 14:10 h light:dark cycle. Fish between 7 weeks and 18 months old were used for experiments. Scale plucking was performed essentially as previously described (*9, 12, 13*). In brief, fish were anaesthetized in 0.075% phenoxyethanol or 0.016% tricaine in system water until swimming ceased and operculum movement slowed. Then, they were placed in a Petri dish, and fluorescent scales were viewed under a fluorescence dissecting microscope. Three rows of 6–8 scales, unless differently indicated, were plucked with forceps from the trunk of the fish, starting from the caudal peduncle and proceeding anteriorly. After scale removal, fish were returned to system water to recover from anesthesia. No statistical methods were used to predetermine sample size. Mixed males and females, when possible siblings, were randomly allocated in control and experimental groups. It was not possible to blind investigators during data collection, because fish from control and experimental groups can be distinguished by cell behaviors or fluorescent reporters. However, data quantification was performed automatically using the same computational algorithm. When necessary, human manual data curation was performed by blinded researchers, although often data from control and experimental groups can be recognized from phenotypes. Researchers were not blinded during data visualization. Fish were euthanized in 0.09% phenoxyethanol or 0.032% tricaine in system water. Fish husbandry and animal experiments were approved by the Service de la Consommation et des Affaires Vétérinaires of the République et Canton de Genève (GEH8A and GE225A-H/GE547, respectively).

### Construction of transgenic zebrafish

The Tg(*osx:p38KTR-mVenus*) fish line was generated by I-SceI-mediated random insertion of an *osx:p38KTR-mVenus* cassette obtained by synthesis and ligation. Synthesis was based on the sequence pENTR-p38KTR-Clover gifted by M. Covert (*35*). The synthesized construct was ligated into a pBluescript SK plasmid modified to contain I-SceI sites flanking the multicloning site and containing a medaka *osx* regulatory sequence (*29*), derived from a *osx:H2A-mCherry* plasmid (*9*). Plasmids were linearized using I-SceI enzyme (final concentration: 0.33 U μl^−1^) for 30 min at 37 °C before injection into AB embryos at the one-cell stage.

The Tg(*fgfr3PromoterKI:Venus-PEST*) fish line was generated by targeted insertion of a *hspl70:Venus-PEST* cassette, gifted by M. Bagnat. The *hspl70:Venus-PEST* cassette was inserted into the upstream locus of the *fgfr3* coding sequence according to a strategy from (*36, 37*), using CRISPR-Cas9-mediated genome editing. In this strategy, the *hsp70l* promoter was used as a minimal promoter to be activated by *fgfr3 cis-*regulatory sequences. A 5’-ctgcggagtctcagactttatgg-3’ gRNA sequence was used, obtaining insertion of the *hsp70l:Venus-PEST* cassette 8740 bp upstream of the start codon of the *fgfr3* genomic coding sequence. The donor plasmid was amplified using the PCR primers FWD: 5‘-gcgattaagttgggtaacgc-3’ and REV: 5’-tccggctcgtatgttgtgtg-3’. 10 ng/μl of donor PCR products, 50 ng/μl of gRNA and 500 ng/μl of Cas9 protein and Phenol red were co-injected together into one-cell-stage zebrafish embryos.

Plasmids and plasmid sequences are available upon request to the corresponding author.

### Immunofluorescence

After euthanasia, fish were fixed overnight in a 4% paraformaldehyde solution and then transferred to PBS. Scales were collected in PBS and permeabilized. Thereafter, scales were incubated with a blocking buffer, incubated with primary antibodies diluted in blocking buffer, washed, incubated with secondary antibodies diluted in blocking buffer and washed. Finally, scales were mounted on slides.

### Live bone staining

Fish were incubated in 0.2% Calcein in system water for 30 min, followed by a washout in system water for 30 minutes, or in 0.01% Alizarin Red S in system water for 20 minutes, followed by a washout in system water for 5 minutes.

### Pharmacological treatments and heat-shock

For pharmacological treatment with the Mek inhibitor PD0325901 or the pan-Fgfr inhibitor BGJ398, fish were immersed in the pharmacological compound diluted to working concentration in fish water from a stock solution in DMSO for the duration of the treatment. Fish were maintained off the aquarium system in the dark in 50 ml system water per fish for the duration of the treatment. The treatment medium was changed roughly every 12 h. Fish were fed before changing treatment medium. In each experiment, control fish (siblings, when possible) were treated with the same concentration of DMSO vehicle as the treated group and imaged during the same day. The correspondence between fish and scales at different time points was determined before imaging by tank labeling, fish similarity, and location in the fish.

Heat-shocks were performed essentially as previously described (*65*). Fish were placed in an aquarium system with recirculating water that raised temperature from 26–28.5°C to 38 °C in 1h and maintained it for 1 h. Water was allowed to cool down for at least 1 h before imaging. *hsp70l:mCherry-2a-fgf20a* fish were heat-shocked at the same time as their control siblings that did not carry the transgene.

### Confocal imaging

*In vivo* confocal imaging was performed as previously described (*9, 12*). A zebrafish was anesthetized in 0.016% tricaine in system water and transferred to a 1% agarose bed in a custom 3D-printed mount made of polyvinyl alcohol (PVA). The caudal fin of the fish was set on a PVA or glass slide bridge to bring the caudal peduncle of the trunk parallel with the platform, and diluted tricaine was placed near the head of the fish. Cooling 1% agarose was applied on the caudal fin, the trunk anterior to the imaged scale, and the platform areas dorsal and ventral to the scale. Then, fish were immersed in diluted tricaine. Gill movements were monitored visually and, when they slowed, system water was applied using a peristaltic pump with silicone tubing (2.4 mm inner and 4.0 mm outer diameter; 10 ml/min flow rate) until regular rhythm was restored. After imaging, a system water flow was applied using the peristaltic pump, until gill movement quickened. Then, the fish was returned to system water. For longitudinal time courses, fish were mounted, imaged and then returned to system water at each time point. For imaging of scales older than 15 dpp, fish were euthanized and scales were collected in PBS, fixed in a 4% paraformaldehyde solution for 1 h, washed in PBS, and mounted on slides using Prolong Gold Antifade Reagent (Figure 3D).

Confocal images were acquired using a Leica Stellaris 5 confocal microscope and LAS X software with a 25× HC FLUOTAR L VISIR lens at 0.75× zoom or 16× HC FLUOTAR L VISIR lens at 0.75× zoom. Fixed scales were imaged with a 63× PLAN APO lens at 0.75× zoom. As scales are often larger than the field of view of our microscopy setup, they were often imaged with multiple overlapping z-stacks (up to 6 with the 16× and 25× objectives, up to 60 with the 63× objective, with variable number of planes) to cover the entire osteoblast tissue. For some experiments (Figures 6C–H, S4F (first time point) and S4G (first time point)) only a stripe including the scale center and the posterior boundary was imaged. Live scale images were acquired at 1,024 × 1,024 resolution (corresponding to pixel sizes of 0.606 μm, 0.950 μm, and 0.240 μm for the 25×, 16×, and 63× objectives, respectively) and with variable z-step size (between 0.606 μm and 0.75 μm for the 25× objective; 0.950 μm for the 16× objective; between 0.240 μm and 0.600 μm for the 63× objective).

Fluorescent proteins and dyes were imaged using the following lasers: DAPI, 405 nm; ErkKTR-mCerulean, 448 nm; Calcein, 494 nm; Venus-hGeminin, 515 nm; p38KTR-mVenus, 515 nm; Venus-PEST, 515 nm; H2A-mCherry, 561 nm; mCherry-zCdt1, 561 nm; Alizarin Red S, 587 nm; Alexa Fluor 594, 590 nm; Alexa Fluor 647, 653 nm. Laser power and gain were adjusted based on the signal strength of each fluorescent protein or dye.

### Image processing

Image processing was performed in two parts. In the first part, images were processed using custom-written MATLAB code, as previously described (*12*), complemented with custom-written Python code. In brief, image stacks were stitched, overlapping neighboring scales were computationally removed, the scale of interest was aligned to the x-y plane, and the tissue of interest was computationally isolated. Differently from (*12*), registration of time points of the same scale was performed manually with custom-written Python software and using scale radii as a reference (when available). For quantification, the tissue of interest was computationally isolated at a later step using nuclei segmentation. In the displayed raw images (except for Figure S4C), the tissue of interest was computationally isolated by manual data curation. H2A-mCherry was used as reference signal for image processing and data curation, when available; otherwise, a combination of Venus-hGeminin and mCherry-zCdt,1 or p38KTR-mVenus, or Venus-PEST were used.

The second part of the image processing pipeline was custom written in Python. Nuclei were segmented by StarDist using the nuclear signal H2A-mCherry, a combination of Venus-hGeminin and mCherry-zCdt1 or DAPI. Segmented cells were assigned to the episquamal (epidermal) or hyposquamal (dermal; the tissue of interest) osteoblast population based on distance from the segmented dermal boundary of the scale, with a manual correction step. For quantification of Erk activity, FUCCI state, p38 activity, anti-p-p38 levels, and *fgfr3* expression in individual cells, nuclear and cytoplasmic signals were calculated using the segmented nuclear mask and a cytoplasmic mask. Cytoplasmic masks were obtained similarly to (*12*), i.e. by dilating the segmented nuclear mask and subtracting the nuclear mask itself. In Figures 5A and 5B, cells having less than 50 in average level of p38KTR-Venus signal intensity were excluded from quantification (average of cytoplasmic and nuclear regions). In images in which both Calcein and Venus-PEST signals were present (Figure 6F and a subset of scales in 6D, 6E, 6G and 6H), the emission spectrum of Calcein partially overlapped with that of Venus. Spectral unmixing was performed by imaging Calcein in a portion of the spectrum not overlapping with Venus and subtracting this latter Calcein image from the Venus image.

Erk activity and p38 activities were calculated as cytoplasmic-to-nuclear signal ratio. Cdt1 and Geminin levels were calculated as nuclear-to-cytoplasmic signal ratio. A cell was classified as Cdt1^+^ when its Cdt1 level exceeded 2 and its Cdt1 nuclear signal exceeded 5 intensity levels; otherwise, it was classified as Cdt1^−^. Geminin classification was performed analogously. *fgfr3* levels were calculated from the *fgfr3* reporter nuclear signal intensity by applying a log(x+1) transform, then clamping to the 5^th^ and 95^th^ percentiles of its distribution. Anti-p-p38 levels were calculated from cytoplasmic cellular signal intensity by applying a log(x+1) transform, then clamping to the 10^th^ and 90^th^ percentiles of its distribution.

Profiles were calculated by averaging the measurement of interest along a 160 μm-wide stripe along the A-P axis, at the dorsoventral position where the scale reaches its maximum anteroposterior width. For visualization purposes, profiles were smoothened using a Savitzky-Golay filter of order 3 and a window length of 7. Fronts were detected manually for maturity, Erk activity, p38 activity, and fgfr3 expression in 2D images roughly along the anterior-posterior axis, separating regions of low and high percentage of Cdt1^+^Gem^+^ cells, Erk activity, p38 activity or *fgfr3* expression. Fronts were detected automatically for bone (Calcein or Alizarin staining) as the anterior-most position that exceeds the threshold set at 20% of the intensity’s maximum. Cell divisions (Figures 3A, 3B, 4A, 4D, S4A and S4B) were detected manually from time courses and binned in 80 μm × 80 μm bins. Mean cell areas were calculated as the inverse of cell density in 80 μm × 80 μm bins; cell area growth rates (Figures 3A and 3B) were calculated as the change in mean cell area divided by the initial mean cell area and the time interval. To remove segmentation errors, the top 1% of bins by division count and the top and bottom 1% by cell area growth rate were excluded per scale. To quantify cell divisions inside and outside Erk waves (Figures S4A and S4B), the wave region was manually selected. Cells were manually tracked using a custom-written Python script (Figures 2B and 2C). Fits were performed using scipy’s “optimize.curve_fit” function. The maturation front position per scale and time bin (Figure 7B) was calculated as the mean over individual markers’ fronts at one time point and as the mean over time points per time bin.

### Sort-Seq – Osteoblast dissociation and sorting

*osx:mCherry-zCdt1 osx:Venus-hGeminin* fish were used for bulk RNA sequencing (Sort-Seq). Scale regeneration was induced by plucking about 50 scales per fish, in 3 rows, from each side of each fish. At 3 dpp, fish were euthanized and transferred to PBS. Scales were collected in PBS on ice. The tissue was pelleted by centrifugation (5 min at 300g) and resuspended in 600 μl of 13 U ml^−1^ Liberase in HBSS and incubated at 37 °C for 15 min. 500 μl of supernatant were removed and quenched with 65 μl sheep serum on ice. The collected supernatant was filtered using 50 μm filters, pelleted (5 min at 600g) and resuspended in 1 ml DMEM with 1% UltraPure BSA. Cell suspensions were then incubated with 2 μg/ml of Hoechst 33342 at 37 °C for 15 min. Before fluorescence activated cell sorting (FACS) and to exclude dead cells, 1 μM of DRAQ7 was added to the cell suspensions. Cells were analyzed and sorted using a MoFlo Astrios Cell Sorter. Hoechst^+^ DRAQ7^−^ cells were kept and sorted into three groups: mCherry^−^zCdt1^+^ Venus-hGeminin^−^, mCherry-zCdt1^−^ Venus-hGeminin^+^, and mCherry-zCdt1^+^ Venus-hGeminin^+^. Cells were collected in RLT Plus lysis buffer.

### Sort-Seq – RNA extraction and bulk sequencing

RNA was extracted using TRI Reagent followed by purification with the RNeasy microKit. RNA concentration and integrity were assessed with a Bioanalyzer. The SMART-Seq mRNA kit was used for reverse transcription and cDNA amplification according to manufacturer’s specifications, starting with 1 ng of total RNA as input. 200 pg of cDNA were used for library preparation using the Nextera XT kit. Library concentration and quality were assessed with the Qubit and Tapestation using a DNA High sensitivity chip. Libraries were sequenced on a NovaSeq 6000 Illumina sequencer for SR100 reads.

### Sort-Seq – Gene expression analyses

STAR software was used to map the sequencing reads on the Ensembl *Danio rerio* genome primary assembly (GRCz11). Gene expression quantification was carried out using SAMtools and htseq. Differential gene expression analysis was performed using DESeq2 software in R. Genes with fewer than 10 counts in at least 4 samples were excluded prior to analysis and size factor normalization was applied to raw counts. Principal component analysis was conducted on the 500 most variable genes following regularized log transformation. Differential expression was assessed by Wald test using a pairwise contrast between the two conditions of interest.

### Single-cell RNA sequencing

Tissue collection and cell dissociation were performed as for Sort-Seq but using *osx:H2A-mEos2* fish. Cell suspensions were incubated with 2 μg/ml of Hoechst 33342 at 37 °C for 15 min. Before fluorescence activated cell sorting (FACS) and to exclude dead cells, 1 μM of DRAQ7 was added to the cell suspensions. After FAC-sorting for H2A-mEos^+^ Hoechst^+^ Draq7^−^ osteoblasts, cells were pelleted by centrifugation (5 min at 1000 rpm), resuspended in 100 μl DMEM, and counted using a Neubauer chamber with Trypan Blue exclusion (1:10 dilution in DMEM). The concentration was adjusted to 1,000–2,000 cells/μl, and up to 40 μl of the cell suspension was placed on ice. Approximatively 16,000 cells were prepared with the 10x Genomics Chromium Next GEM Single Cell 3’kit v3.1 for loading on the Chromium controller, according to the manufacturer’s instructions. cDNA was quality controlled using Bioanalyzer. Library concentration and quality were assessed with the Qubit and TapeStation. Libraries were sequenced on a NovaSeq 6000 sequencer for PE28-90 reads. FASTQ files were used as input to the 10x Genomics Cell Ranger pipeline.

### Single-cell RNA sequencing - Gene expression analyses

FASTQ files were pre-processed with Cell Ranger with default settings. Reads were mapped on the *Danio rerio* genome primary assembly (GRCz11) using the STAR aligner software implemented in Cell Ranger. The filtered feature-barcode matrix was used for downstream analysis. This matrix included a total of 10,013 cells. The count matrix was then analyzed with Seurat software in R. Cells were retained if they had between 200 and 2,500 detected genes and less than 5% mitochondrial read content; genes expressed in fewer than 3 cells were also excluded. Counts were log-normalized (scale factor 10,000), and the 2,000 most variable genes were identified by variance-stabilizing transformation for downstream dimensionality reduction. Data were scaled across all genes prior to PCA, and a nearest-neighbor graph was constructed on the top 20 principal components to generate UMAP embeddings. Unsupervised clustering was performed using the Louvain algorithm and cluster marker genes were identified with Seurat’s “FindMarkers” function. Cell cycle phase was assigned using Seurat’s CellCycleScoring function, based on canonical S-phase and G2/M-phase gene sets (Tirosh et al., 2016). As the default gene lists are human, we substituted zebrafish orthologs identified via ZFIN. Single cells were assigned to FUCCI-sorted population identities (Cdt1^+^ Geminin^−^, Cdt1^−^ Geminin^+^, Cdt1^+^ Geminin^+^) based on the highest Pearson correlation between their scaled expression profiles and log2 fold changes from bulk RNA-seq comparisons of the sorted populations.

## Acknowledgments

We thank Viviane Pilloux for assistance with experiments and data curation; Thomas Ferrez and Pietro Albanito for data curation; Sandra Gitto, Valérie Haechler and Amelie Waldmann for technical support; Denis Benoni, Julie Zuppiger, Thomas Ferrez, Melanie Dalzell and Yasmina Rafrafi for fish maintenance; the Flow Cytometry core facility of the University of Geneva for assistance with FACS experiments; the iGE3 Genomics platform of the University of Geneva for assistance with bulk and single-cell RNA-seq experiments; Marcos Gonzalez Gaitan, Tristan Guyomar and Pierre Gönczy for critical reading of the manuscript. We are grateful to M. Covert and M. Bagnat for sharing plasmids.

## Funding

This work has received funding from:

The Swiss National Science Foundation under Eccellenza Professorial Fellowship (PCEFP3_202776; ADS).

The Swiss National Science Foundation under SNSF project grant (200021_197068; GS).

The Swiss State Secretariat for Education, Research and Innovation (SERI - Transitional Measure MB22.00070; ADS).

## Author contributions

KM conceived the project, designed the experiments, performed experiments, developed the data analysis software, curated data, analyzed data, visualized data, contributed to developing the theory and wrote the initial draft. SC contributed to conceiving the project, contributed to designing experiments, developed the theory and visualized theory results. DC performed experiments and curated data. GS contributed to conceiving the project, contributed to designing experiments, developed the theory, provided supervision and acquired funding. ADS conceived the project, designed the experiments, analyzed data, contributed to developing the theory, wrote the initial draft of the manuscript, acquired funding and provided supervision.

All authors reviewed and edited the final draft.

