## Supplemental Materials for "A moving front of osteoblast maturation scales regenerating zebrafish bone"

Konrad Marx *et al.*

**This PDF file includes:**

Figures S1 to S4

Supplementary Information

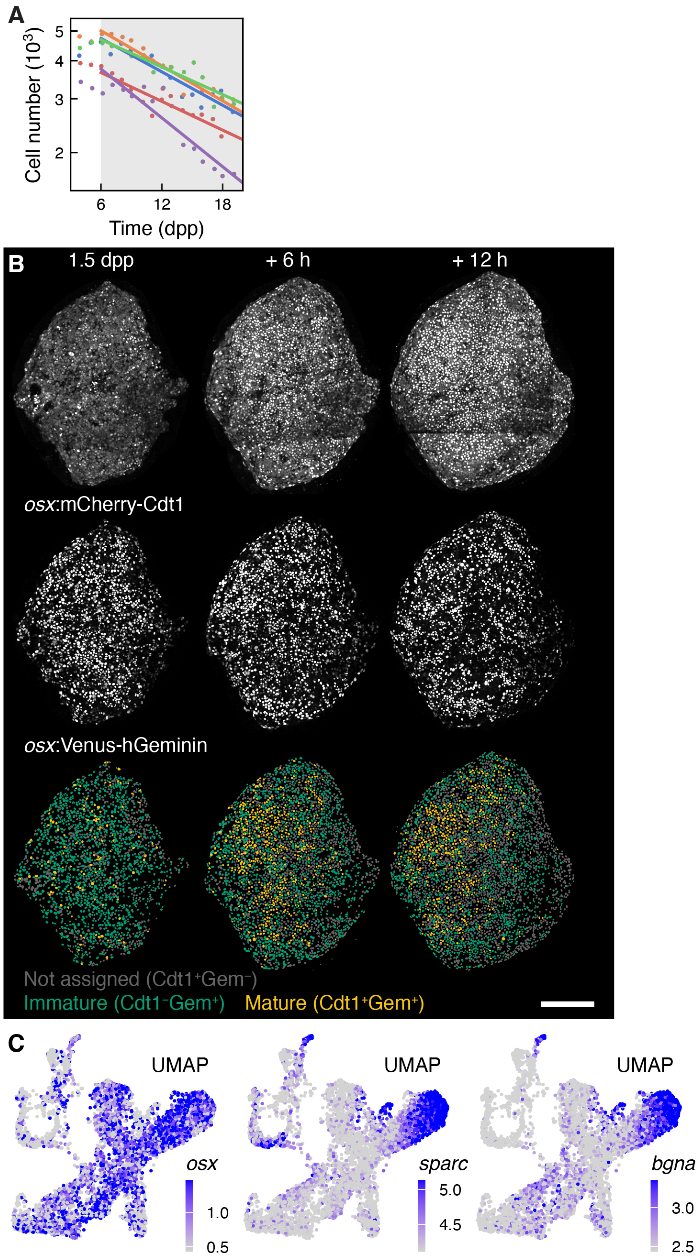


**Figure S1. Osteoblast number dynamics in late scale regeneration. Raw Cdt1 and Gem signals with quantification and dynamics as a function of cell-cycle time. Single-cell expression of *osx*, *sparc,* and *bgna* transcripts.**

A. Cell number over time $N(t)$ during late stage of regeneration in individual scales (with linear least-squares fit of $\ln N(t)$ after 6 dpp (shaded area), mean slope $-0.0018$ $h^{-1}$; n = 5 scales from 5 fish, from 1 experiment re-analyzed from [13]). **B**. Example of Cdt1 and Geminin signals, assignment to $Cdt1^{\pm}$ $Geminin^{\pm}$ populations. **C**. Visualization of *osx* (left), *sparc* (center) and *bgna* (right) transcripts per cell in scSeq experiment. Color bar: Seurat log-normalized transcript counts. Scale bar, 200 μm. Dpp: days post-plucking. Gem: Geminin.

A

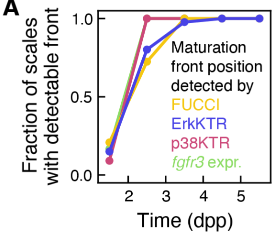


**Figure S2. A front of osteoblast maturation and signaling changes is established between 1.5 and 2.5 dpp.**

**A**. Fraction of scales with detectable front. Maturity front (FUCCI; see Figures 2L, 2M, 3E, 3F and S3A): n = 35 scales from 16 fish, from 8 experiments; Erk activity (ErkKTR; see Figures 4A–D, 4I, 4J, 5C, 5E and S3A): n = 85 scales from 41 fish, from 20 experiments; p38 activity (p38KTR; see Figures 5C and 5D): n = 18 scales from 9 fish, from 4 experiments; *fgfr3* expression (see Figures 4E–J): n = 24 scales from 9 fish, from 3 experiments; a front of stained mineralized bone (Calcein or Alizarin) could be detected at all time points (see Figures 3C–F, 4G and 4H): n = 74 scales from 31 fish, from 12 experiments. Act: activity. Expr: expression. Dpp: days post-plucking.

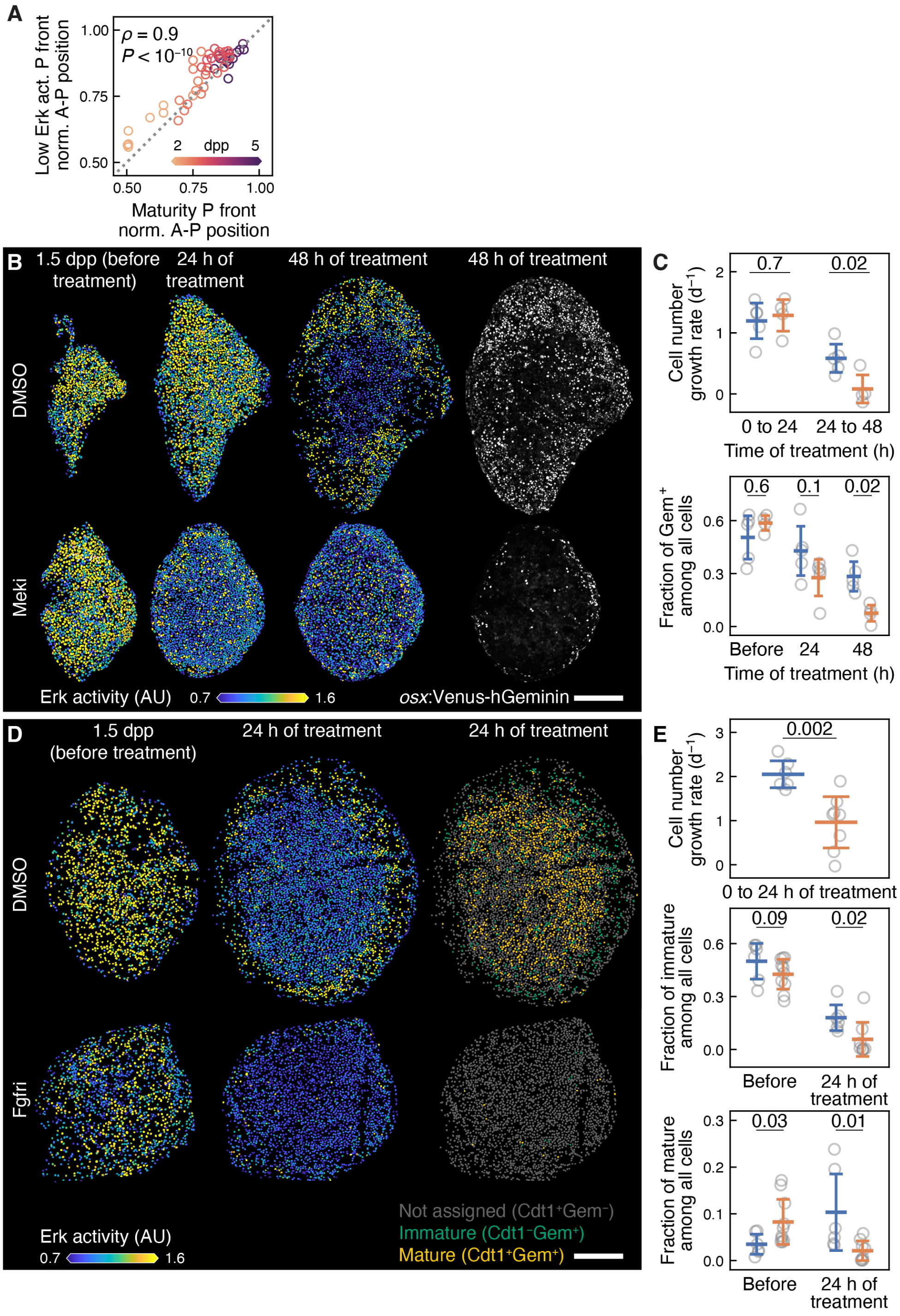


**Figure S3. Erk activation is required for osteoblast proliferation in zebrafish scales.**

**A**. Position of the posterior front of the low Erk activity region as a function of the position of the posterior front of the mature Cdt1$^+$ Geminin$^+$ region (circles: individual scales and time points from n = 35 scales from 16 fish; from 8 experiments; Pearson's correlation coefficient ρ with P-value is indicated; dashed line: bisector of the axis). **B**. Example of Erk activity (left) and Geminin signal (right) in control (DMSO) and treated (Mek inhibitor PD0325901, 10 μM) fish. **C**. Cell number growth rate and fraction Geminin$^+$ cells in control (DMSO) and treated

(Mek inhibitor PD0325901, 10 µM) fish (circles: individual scales; DMSO n = 5 scales from 5 fish, Meki n = 4–5 scales from 3–4 fish; from 2 experiments; with SD). Student's unpaired t-tests' (cell number growth rate) and Mann-Whitney U tests' (fraction of $Gem^+$ cells) P-values are indicated. **D**. Example of Erk activity and $Cdt1^\pm$ $Geminin^\pm$ assignment in control (DMSO) and treated (pan-Fgfr inhibitor BGJ398, 10 µM) fish. Scale bars, 200 µm. AU: arbitrary units. **E**. Cell number growth rate, fraction of immature $Cdt1^-$ $Geminin^+$ cells and fraction of mature $Cdt1^+$ $Geminin^+$ cells in control (DMSO) and treated (pan-Fgfr inhibitor BGJ398, 10 µM) fish (circles: individual scales; DMSO n = 6 scales from 3 fish, Fgfri n = 8–10 scales from 4–5 fish; from 1 experiment). Student's unpaired t-test's (cell number growth rate) and Mann-Whitney U tests' (fractions) P-values are indicated. Dpp: days post-plucking. Gem: Geminin.

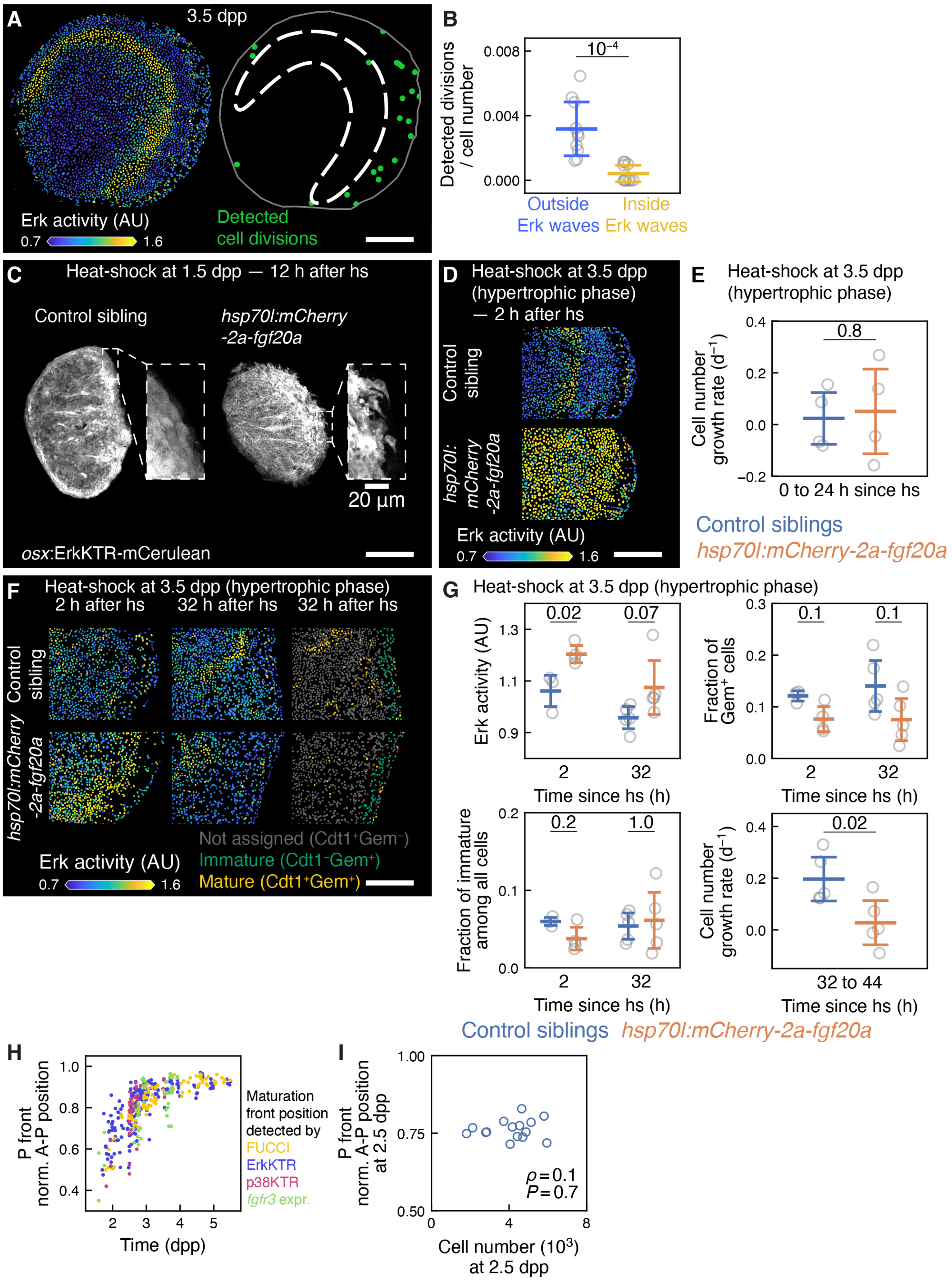


**Figure S4. Example of osteoblast behaviour after Fgf over-expression. Erk activation is not sufficient to trigger cell proliferation once osteoblasts have matured. Maturation front.**

**A**. Example of Erk activity and distribution of detected cell divisions at the onset of the hypertrophic phase. Dashed line: Erk activity wave. **B**. Detected cell divisions per cell inside and outside the Erk activity wave region (circles: individual scales; n = 4 scales from 2 fish; from 1 experiment; with SD; Student's paired test's P-value is indicated). **C**. Example of osteoblasts migrating away from the scale following heat-shock performed at 1.5 dpp in *hsp70l:mCherry-2a-fgf20a* fish, compared with control sibling, visualized using the ErkKTR signal. **D**. Example of Erk activity after heat-shock at 3.5 dpp in *hsp70l:mCherry-2a-fgf20a* and control fish. **E**. Cell number growth rate at different times following heat-shock at 3.5 dpp in *hsp70l:mCherry-2a-fgf20a* and control fish (circles: individual scales; control n = 4 scales from 2 fish, *hsp* n = 4 scales from 2 fish; from 1 experiment; with SD; Student's unpaired t-test's P-value is indicated). **F**. Example of Erk activity and mature and immature cells after heat-shock at 3.5 dpp in *hsp70l:mCherry-2a-fgf20a* and control fish. **G**. Erk activity, fraction of Geminin+ cells, fraction of immature cells and cell number growth rate at different times following heat-shock at 3.5 dpp in *hsp70l:mCherry-2a-fgf20a* and control fish (circles: individual scales; control n = 3–5 scales from 2–3 fish, *hsp* n = 4–5 scales from 2–3 fish; from 1 experiment; with SD; Student's unpaired t-tests' (Erk activity and proliferation) and Mann-Whitney U tests' (fractions) P-values are indicated. **H**. Maturation front A-P position over time (dots: fronts from individual scales and time-points; data from Figures 2M, 4B, 4F and 5D). **I**. Size of the maturation front at 2.5 dpp (circles: individual scales; Pearson's correlation coefficient ρ with P-value is indicated; from Figure 7A). Scale bars, 200 µm, unless differently indicated. Act: activity. A-P: anteroposterior axis. AU: arbitrary units. Dpp: days post-plucking. Expr: expression. Gem: Geminin. Norm: normalized.

**Supplementary Information**

*Mathematical model of proliferation dynamics*

Assuming radial symmetry, we describe the evolution of the maturation front in terms of a relative radial coordinate $r(t) = R_{\text{front}}(t)/R_{\text{scale}}(t)$, where $R_{\text{front}}(t)$ and $R_{\text{scale}}(t)$ are the radii of the mature region and the scale at time $t$, respectively. The time evolution of the number of osteoblasts $N$ reads

$$\frac{dN}{dt} = k_d N_p(t) - \beta N(t)\,, \tag{1}$$

where $k_d$ is the cell division rate, assumed constant for simplicity, $N_p(t)$ is the number of proliferative cells at time $t$, and $\beta$ is the rate of cell death. After the time $t_0$ of appearance of the maturation front the number of proliferative cells is given by

$$N_p(t) = n\left(\pi R_{\text{scale}}^2(t) - \pi R_{\text{front}}^2(t)\right)\,, \tag{2}$$

where $n$ is the cell density, assumed to be uniform and constant for simplicity. In terms of the total number of cells $N(t)$, the equation governing the cell number evolution becomes

$$\frac{dN}{dt} = \left[k_d\left(1 - r^2(t)\right) - \beta\right]N(t)\,, \tag{3}$$

using that $N = n\pi R_{\text{scale}}^2(t)$. Now, we impose the dynamics of the front observed in experiments (Figure 7A). The position of the maturation front, taken from the anterior edge and normalized to the anterior-posterior tissue length, denoted $x_F$, has a dynamics which is well described by an exponential relaxation:

$$x_F(t) = x_{\text{F, max}} + \left(x_{\text{F,0}} - x_{\text{F,max}}\right)e^{-\frac{t-t0}{\tau}} \quad \text{for } t > t_0\,, \tag{4}$$

with $t_0 = 48$ hpp. Since $x_F = (R_{\text{front}} + R_{\text{scale}})/(2R_{\text{scale}})$, we have $r(t) = 2x_F(t) - 1$ and the equation for cell number growth becomes

$$\frac{dN}{dt} = \left[4k_d x_F(t)\left(1 - x_F(t)\right) - \beta\right]N(t) \quad \text{for } t > t_0, \tag{5}$$

with $x_F(t)$ given by Equation (4). Given the initial condition $N(t_0) = N_0$, this equation admits the analytical solution

$$N(t) = N_0 \exp\{\beta(t_0 - t) - 4k_d(t - t_0)(x_{\mathrm{F,max}} - 1)x_{\mathrm{F,max}} \\ +2k_d\tau(x_{\mathrm{F},0} - x_{\mathrm{F,max}})[e^{-\frac{2(t-t_0)}{\tau}}(x_{\mathrm{F},0} - x_{\mathrm{F,max}}) \\ +2e^{-\frac{(t-t_0)}{\tau}}(2x_{\mathrm{F,max}} - 1) - (x_{\mathrm{F},0} - 2 + 3x_{\mathrm{F,max}})]\}, \tag{6}$$

which is plotted in Figure 7D (right). In Figure 7E (left) we plot the time derivative of the cell number at $t_0$,

$$\frac{dN(t_0)}{dt} = N_0[4k_d x_{\mathrm{F},0}(1 - x_{\mathrm{F},0}) - \beta], \tag{7}$$

and the cell number at 6 dpp $N(144\ \mathrm{hpp})$ (right) against the initial cell number $N_0$.

*Parameters*

We use the measured values $x_{\mathrm{F},0} = 0.639$, $x_{\mathrm{F,max}} = 0.95$, $t_0 = 48$ hpp, and $\tau = 24$ h (Figure 7A). The rates of cell division $k_d$ and cell death $\beta$ are estimated from the observed $\frac{1}{N}\frac{dN}{dt}$ calculated before the appearance of the front $(t < 48\ \mathrm{hpp})$ and when the front reached a plateau $(t > 144\ \mathrm{hpp})$. $\frac{1}{N}\frac{dN}{dt}(\mathrm{t} < 48\ \mathrm{hpp}) = 0.049\ \mathrm{h}^{-1}$ is calculated as $\frac{1}{N}\frac{N(t_2)-N(t_1)}{t_2-t_1}$ from experimental data of $N(t)$ with $t_1, t_2$ values such that $t_1 < 42$ hpp and $t_2 - t_1 < 6$ h (n = 31 scales from 15 fish, from 9 experiments). $\frac{1}{N}\frac{dN}{dt}(\mathrm{t} > 144\ \mathrm{hpp}) = -0.0018\mathrm{h}^{-1}$ was calculated as the slope of a linear least-squares fit of $\ln N(t)$, performed independently for each scale (from Figure S1A; n = 5 scales from 5 fish, from 1 experiment re-analyzed from [13]). Before the maturation front emerges, the equation for the evolution of cell number is simply $\frac{dN}{dt} = (k_d - \beta)N$. Therefore, by solving the system of equations

$$\frac{1}{N}\frac{dN}{dt}(\mathrm{t} < 48\ \mathrm{hpp}) = k_d - \beta, \tag{8}$$

$$\frac{1}{N}\frac{dN}{dt}(\mathrm{t} > 144\ \mathrm{hpp}) = 4k_d\, x_{\mathrm{F,max}}(1 - x_{\mathrm{F,max}}) - \beta, \tag{9}$$

we obtain the values $k_d = 0.063\ \mathrm{h}^{-1}$, $\beta = 0.014\ \mathrm{h}^{-1}$.
